# Development and applications of sialoglycan-recognizing probes (SGRPs) with defined specificities: exploring the dynamic mammalian sialoglycome

**DOI:** 10.1101/2021.05.28.446202

**Authors:** Saurabh Srivastava, Andrea Verhagen, Aniruddha Sasmal, Brian R. Wasik, Sandra Diaz, Hai Yu, Barbara A. Bensing, Naazneen Khan, Zahra Khedri, Patrick Secrest, Paul Sullam, Nissi Varki, Xi Chen, Colin R. Parrish, Ajit Varki

## Abstract

Glycans that are abundantly displayed on vertebrate cell surface and secreted molecules are often capped with terminal sialic acids (Sias). These diverse 9-carbon-backbone monosaccharides are involved in numerous intrinsic biological processes. They also interact with commensals and pathogens, while undergoing dynamic changes in time and space, often influenced by environmental conditions. However, most of this sialoglycan complexity and variation remains poorly characterized by conventional techniques, which often tend to destroy or overlook crucial aspects of Sia diversity and/or fail to elucidate native structures in biological systems i.e., in the intact sialome. To date, *in situ* detection and analysis of sialoglycans has largely relied on the use of plant lectins, sialidases or antibodies, whose preferences (with certain exceptions) are limited and/or uncertain. We took advantage of naturally-evolved microbial molecules (bacterial adhesins, toxin subunits and viral hemagglutinin-esterases) that recognize sialoglycans with defined specificity to delineate 9 classes of Sialoglycan Recognizing Probes (SGRPs: SGRP1–SGRP9) that can be used to explore mammalian sialome changes in a simple and systematic manner, using techniques common in most laboratories. SGRP candidates with specificity defined by sialoglycan microarray studies were engineered as tagged probes, each with a corresponding non-binding mutant probe as a simple and reliable negative control. The optimized panel of SGRPs can be used in methods commonly available in most bioscience labs, such as ELISA, Western Blot, flow cytometry and histochemistry. To demonstrate the utility of this approach, we provide examples of sialoglycome differences in tissues from C57BL/6 wild type mice and human-like *Cmah*^−/−^ mice.

## Introduction

All cells in nature are covered with a dense and complex array of sugar chains (1), and in vertebrates the outermost ends of the branches on this glycan forest are often capped with monosaccharides called sialic acids (Sias), which have enormous intrinsic complexity (2, 3). Most current methods to study this important and dynamic aspect of the glycome are too specialized for an average scientist to employ, and many aspects of this class of important molecules are thus poorly studied, even by experts. Given their ubiquitous presence and terminal position, Sias have been exploited as primary, transient or co-receptors by a diverse range of commensal or pathogenic microorganisms (4, 5). These interactions are typically mediated by microbial proteins that have evolved with high binding specificity towards sialoglycans and can differentiate their target Sias by types, modifications, substitutions and/or linkage to underlying glycans.

In this report, we undertake a strategy to develop Sia-specific probes from such microbial proteins that have naturally evolved to recognize Sia complexity with great specificity, likely because of the ongoing evolutionary arms race between microbes and hosts. We have harnessed the Sia specificity of microbial proteins, and if found insufficient assessed other available probes to generate a simple and reliable toolkit that can be used to easily monitor dynamic changes of the sialic acids in normal and abnormal states. Specifically, this study advances a set of comprehensive sialoglycan-recognizing probes (SGRPs) that can confirm whether a biological sample has any sialic acids or not; if so, the type of common Sia variations (particularly O-acetylation), the linkage to underlying glycans, and the presence of *N*-acetyl or *N*-glycolyl groups. To identify candidates for such an updated set of probes for mammalian Sias, we assessed a number of Sia-specific binding proteins, novel or reported in accompanying or previous articles for their specificities towards different classes of mammalian Sia types and/or linkage to the underlying glycans.

Specifically, 9 types of SGRPs were defined from a subset of proteins including bacterial serine-rich repeat (SRR) adhesins, bacterial B5 toxins, viral hemagglutinins (HAs) and hemagglutinin-esterases (HEs) and compared them to previously known invertebrate and plant lectins, selected Siglecs, and polyclonal and monoclonal antibodies (MAbs). Upon identification of the best SGRP for each class of sialoglycans, a mutant inactive probe was also developed as an internal control of each probe’s specificity. To ensure minimal loss of sensitive Sia modifications/substitutions, experimental conditions were optimized, and the specificity of each probe was tested with positive and negative controls, such as pretreatment with specific sialidases or esterases, or mild periodate oxidation of the Sia side chain. The binding specificities of SGRPs were confirmed by testing on a sialoglycan microarray, constituting a diverse array of more than 100 mammalian sialoglycans and demonstrated by examples of generalized laboratory methods of ELISA, Western blotting, and fluorescence detection by flow cytometry and histological analysis. An example of application of SGRPs is provided, showing sialoglycome changes in mice with human-like loss of the CMAH enzyme.

## Results and Discussion

### Defining distinct classes of Sialoglycan Recognizing Probes (SGRPs)

We sought to define a set of SGRPs for detection of the most common types of mammalian sialoglycan variants, along with non-binding mutants as controls. This approach simplifies *in-situ* detection for all major types of Sias (SGRP1), typical *N*-acyl modifications at C-5 of Sias (SGRP2, SGRP5), linkages to underlying glycans (SGRP3, SGRP6, SGRP8) and occurrence of *O*-acetyl groups (SGRP4, SGRP7, SGRP9). We also suggested a SGRP numbering system (see Table 1) for these probes that make them easier to remember. To define a given probe, we used a superscript, and added NB for the non-binding control. Thus, for example, the *Y. enterocolitica* toxin B subunit (YenB) that recognizes all Neu5Ac and Neu5Gc forms and linkages is designated SGRP1*^YenB^* and the nonbinding variant as SGRP1*^YenB^*NB. The current set of SGRPs does not include probes specific for 2-keto-3-deoxy-D-glycero-D-galacto-nononic acid (Kdn), which, although naturally found in mammals, occurs in limited amounts and primarily in the free form (6).

**Table 1.**
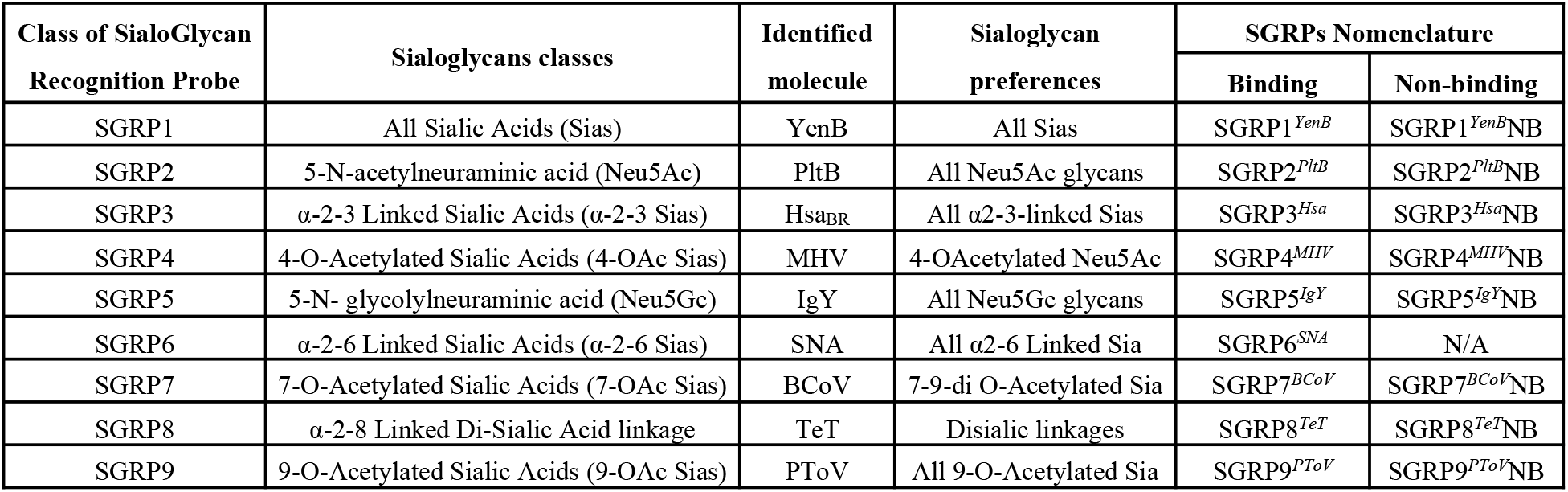
Sialoglycan Recognition Probes and their specificities. Table representing the classes of Sialoglycan Recognition Probes (SGRPs), their presumed binding specificities, the most appropriate molecules as observed by assessment of sialoglycan binding by various glycomic methods, their experimentally confirmed sialoglycan preferences, and defining SGRP nomenclature suggesting class of probe along with source of probe as mentioned in superscripts. The names of probes ending with NB represent the non-binding variant of SGRPs.

### Search for naturally evolved microbial molecules with defined specificity towards specific aspects of sialoglycans

The SGRP candidates were defined by sialoglycan microarray studies, with each probe accompanied by a corresponding non-binding mutant as a negative control. The essential criterion for a protein to qualify as an SGRP was that it must show specificity towards the preferred Sia modification/linkages. The detailed analyses of most updated contemporary set of probes (SGRPs) and specificities and their practicability towards probing mammalian sialoglycans is discussed in subsequent sections. As mentioned in Table 1, we could not identify any microbial candidate having a better or broader specificity for SGRP5 (*N*-glycolyl-Sias) than our previously described affinity--purified Neu5Gc chicken polyclonal IgY, or SGRP6 *Sambucus nigra* agglutinin (SNA), the conventionally used lectin probe for α2-6-linked Sias.

### SGRP1*^YenB^* is a probe to detect all mammalian sialic acids

While α2-3Sia-binding MAL and α2-6Sia-binding SNA together recognize the majority of Sia linkages, their preferences cannot be generalized for all types of Sias. Previously, WGA and LFA have been reported as having broad-spectrum Sia specificity, but their preferences towards Neu5Ac bring into question their utility as practicable probes to detect all types of Sias (7–9). For an SGRP that binds all types of Sia, we first considered YpeB (Yersinia pestis Toxin B subunit that recognizes both Neu5Ac- and Neu5Gc-terminated glycans––detailed sialoglycan preferences are reported in the companion paper 1 (Khan et al., see also Figure S1). However, despite its ability to recognize most major classes of mammalian sialoglycans in glycan array and serum ELISAs, YpeB did not bind 4-OAc-Sias (Suppl. Fig S1). Relying on the promising lead from YpeB, we investigated additional B subunits of AB5 bacterial toxins and selected YenB (*Y. enterocolitica* toxin B subunit) based on homology with YpeB and *S. typhimurium* ArtB, and broad host specificity (see companion paper, Sasmal et.al.). Using His6-tagged YenB (see Sasmal et al. accompanying paper) we confirmed the protein’s recognition for both Neu5Ac and Neu5Gc including 9-*O*- and 4-*O*-acetylated Sias, a clear advantage over WGA, LFA and our initial candidate YpeB (Suppl. Figure S2). Notably, YenB did not show any binding to non-sialylated glycans in the same assay, further confirming its extensive but Sia-selective binding to glycans. In order to biotinylate YenB without affecting its Sia-binding domain, we attempted to clone YenB with additional tags for biotinylation (SNAP, ACP and Avi) but the modifications resulted in poor protein quality and yield, leading to reduced binding of Sia in glycan arrays. A direct *N*-hydroxysuccinimide (NHS-)-biotin conjugation of YenB was therefore optimized to obtain the final biotinylated probe (SGRP1*^YenB^*) that showed no loss of binding preferences in comparison to non-biotinylated YenB. It was previously established that a serine residue contributes critically to Neu5Ac binding, while a tyrosine residue interacts with the extra OH group at the C5-acyl chain of Neu5Gc and is thus critical for its binding (10). As described in a companion paper (Sasmal et.al.), we aligned the YenB sequence with those of *E. coli* SubB and *S. typhimurium* ArtB and predicted the conserved serine (S31) and tyrosine (Y100) in YenB. Mutating these critical sites for Sia recognition (YenB, S31A;Y100F) produced SGRP1*^YenB^*NB as an internal control for Sia-binding SGRP1*^YenB^* (Suppl. Fig. S3A &B). The final binding and nonbinding specificities for SGRP1*^YenB^* were tested on a sialoglycan microarray with nearly 130 mammalian sialoglycan types, suggesting all-inclusive Sia specificity in SGRP1, and a complete lack of Sia-binding by SGRP1*^YenB^*NB (Suppl. Fig S4). Since there were no other molecules known to possess YenB-like Sia specificities, the pair of SGRP1*^YenB^* and SGRP1*^YenB^*NB are currently the most appropriate probes to detect all mammalian Sia types. The utility of SGRP1*^YenB^* as a useful tool of *in situ* Sia detection through ELISAs, Western Blotting, IHC and flow cytometry is described and discussed below.

### SGRP2*^PltB^* recognizes all Neu5Ac-terminated glycans

Neu5Ac is the most abundant Sia type and occurs in all Sia-expressing organisms. Mammals (including humans) that lack a functional CMAH enzyme have lost the ability to convert cytidine 5’-monophosphate *N*-acetylneuraminic acid (CMP-Neu5Ac) into CMP-Neu5Gc, and express excess Neu5Ac as their primary Sia type. Despite such wide distribution and roles in human physiology/immunity, there has so far been no direct probe to selectively detect all forms and linkages of Neu5Ac *in situ*. Among commonly known probes for sialoglycans, WGA shows relative specificity towards Neu5Ac-glycans but neither recognizes all sialoglycans terminating with Neu5Ac, nor is it exclusive to Neu5Ac, showing a dual preference for Neu5Ac and GlcNAc (8) (Suppl Fig. S2A). Previously, we identified Neu5Ac-specific binding in PltB, the B subunit of Typhoid toxin which preferred to bind human erythrocytes and tissues rich in Neu5Ac over the corresponding Neu5Gc-rich samples from Chimpanzees (11). Similar Neu5Ac-specific patterns of PltB binding were further studied recently by others who also reported its binding *O*-acetylated Neu5Ac in both α2-3 and α2-6-linked sialoglycans (12). In the accompanying paper (Sasmal et al.), we report on the binding of PltB to a range of naturally occurring Neu5Ac but not Neu5Gc-glycans, underlining PltB’s appropriateness as a Neu5Ac-recognizing probe. We cloned the His6-tagged Neu5Ac-binding domain of PltB with an additional ACP tag (NEB) for biotinylation, but we then had to abandon this approach due to poor expression and purification quality of the resulting protein. The final probe version of PltB (SGRP2*^PltB^*) was derived by direct biotinylation of PltB with NHS-biotin. This biotinylated-PltB shows comparable specificity and avidity towards Neu5Ac glycans as the unmodified protein (Fig 1). The Neu5Ac-binding domain of PltB was characterized previously and a serine residue was reported to be critical for Neu5Ac-binding by PltB (11). Thus PltB^S35A^NB, an internal nonbinding mutant control of SGRP2 was also produced, biotinylated and included in all studies (Suppl. Fig S3 C & D).

**Figure 1.**
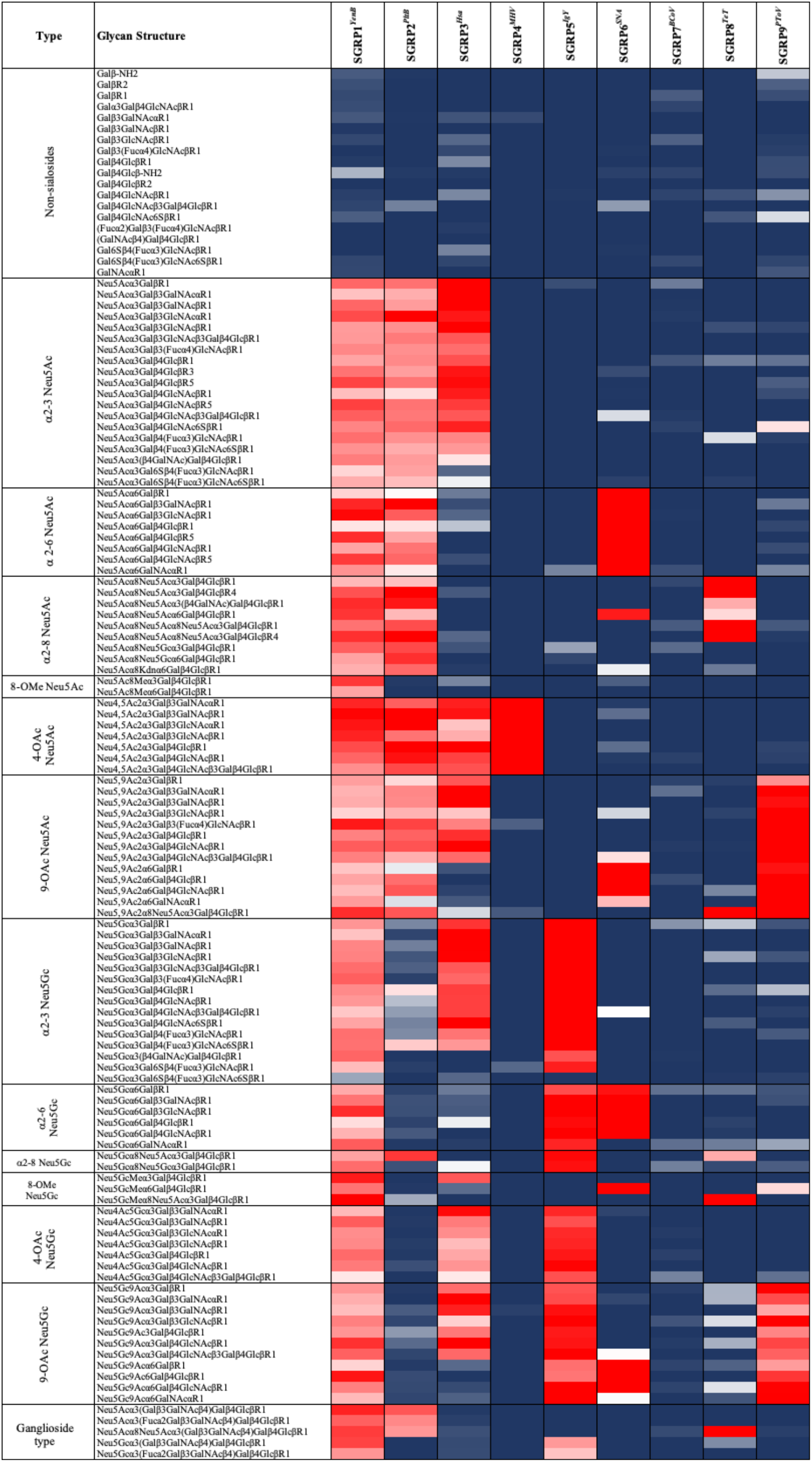
Sialoglycan Microarray Binding Studies of Proposed Sialoglycan Recognizing Probes (SGRPs). Heatmap analysis of SGRPs (see Table 1 for details of nomenclature) binding to mammalian (sialo) glycans; SGRP1*^YenB^* (30 μg/ml), SGRP2*^PltB^* (30 μg/ml), SGRP3^Hsa^ (30 μg/ml), SGRP4*^MHV^* (30 μg/ml), SGRP5*^IgY^* (20 μg/ml), SGRP6*^SNA^* (20 μg/ml), SGRP7*^BCoV^* (60 μg/ml), SGRP8*^TeT^* (30 μg/ml) and SGRP9*^PToV^* (30 μg/ml) in microarray experiments. SGRPs binding efficiencies are displayed in red for highest binding (saturated or 100%), blue for minimum or no binding (0%) and intermediate binding represented by colors ranging between blue and red. Ranks; Red (100%, or maximum), Blue (0%, or minimum). Markedly reduced or absent binding was seen with the no-binding control probes studied simultaneously (See Supplementary Figure S4).

The sialoglycan microarray with mammalian sialoglycan types showed the Neu5Ac-specific binding pattern of SGRP2*^PltB^* while SGRP2*^PltB^*NB completely lacks binding (Suppl. Fig S4). In agreement with previous data SGRP*^PltB^* (11), high binding preferences towards Neu5Ac were seen regardless of substitutions, modifications, or linkages to underlying glycans. As shown in the array, SGRP2*^PltB^* demonstrates comparable binding for α2-3, α2-6 and α2-8 linked Neu5Ac and clearly distinguishes them from α2-3, α2-6 or α2-8 linked Neu5Gc (Fig 1). Notably, SGRP2*^PltB^* also exhibited recognition for Neu4,5Ac2 and Neu5,9Ac2. Taken together, SGRP2*^PltB^* showed definite superiority over WGA as Neu5Ac binding probe (Suppl Fig S2A). We therefore prepared SGRP2*^PltB^* and a SGRP2*^PltB^*NB control as Neu5Ac-binding and non-binding probes for use in general methods such as ELISAs, Western blotting, flow cytometry and IHC.

### SGRP3*^Hsa^* selectively recognizes α2-3-linked Sias

Lectins from *Maackia amurensis* seeds, first reported by Kawaguchi et al (13) have been the gold standard for detection for Sias α2-3-linked to penultimate Gal. These lectins initially designated as ‘strongly mitogenic *M. amurensis* leukoagglutinin (MAL) and ‘strongly hemagglutinating *M. amurensis* hemagglutinin’ (MAH) are commonly known as MAL-I and MAL-II, respectively. While both lectins require at least a Siaα2-3Gal disaccharide structure to bind, they differ in their requirement for specific structures. MAL-I shows stronger binding to Siaα2-3Galβ1-4GlcNAc/Glc (a trisaccharide common in N-glycans) and MAH (MAL-II) is selective towards Siaα2-3Galβ1-3GalNAc, typically found in O-linked glycans. Among Siaα2-3-linked structures tested, MAH prefers 9-*O*-acetyl Sias with Neu5Ac over Neu5Gc and Ser/Thr-linked O-glycan structures (14–16). However, MAL-I can also recognize 3-*O*-sulfated Gal terminated oligosaccharides i.e., it does not show exclusivity towards sialylated sequences (17). Instead, MAL-I displays widespread affinity towards α2-3-linked Neu5Ac and Neu5Gc, but with selective inclination towards Asn-linked over Ser/Thr-linked glycans. Attempts had been made to improve *Maackia* lectins (18) but this did not yield a general-purpose Siaα2-3-recognizing probe.

We looked into the Sia-binding properties of previously reported Siglec-like-domain-containing ligand-binding regions (BRs) of *Streptococcus gordonii* SRR adhesins (19–21). In particular, we selected GspB-BR of *S. gordonii* strain M99, Hsa-BR of *S. gordonii* Stain DL1 (Challis), and UB10712-BR of *S. gordonii* strain UB10712, and investigated their suitability for a comprehensive Siaα2-3-linkage identifying probe. All 3 BRs were expressed as SNAPf-His6 fusions in the pGEX-3X vector in a bacterial expression system along with their non-binding variants HsaBR (R340E), GspBBR (R484E) and UB10712BR (R338E) (Suppl Fig S3).

The pairs of HsaBR/HsaBRNB, GspBBR/GspBBRNB and UB10712BR/ UB10712BRNB were biotinylated using SNAP-benzyl guanine chemistry and tested for their sialoglycan binding on our microarray. Despite their structural similarities, the 3 biotinylated BRs displayed uniquely different ligand binding profiles, including differential recognition of sialyl Lewis antigens and sulfated glycans. While GspBBR selectively binds sialyl-T antigen (Neu5Acα2-3Galβ1-3GalNAc) and related structures, HsaBR displays broader specificity covering NeuAcα2-3Galβ1-4GlcNAc and sialyl-T antigen (Fig 1 & Suppl Fig S5A) (20–22). In comparison to HsaBR, GspBBR imparts lesser specificity towards α2-3 than α2-6 Sia linkages, (20) and falls short of the binding range of HsaBR, which not only includes trisaccharide and oligosaccharide but also disaccharide Sias. 9-*O*-Acetylation on Sia did not block GspBBR or HsaBR binding, but sulfation appeared to enhance HsaBR binding (21). Sia binding preferences of UB10712BR remained comparable to α2-3 linked Sia specificities of HsaBR and GspBBR. UB10712BR bound to range of α2-3 Sia linkages including sialyl Lewis X, 3’-sialyllactosamine and their sulfated forms but preferred Neu5Ac over Neu5Gc sequences generally (Suppl Fig S5A) (19)). Confirming the Sia-specificity of these BRs, the NB variants did not show any binding of Sia linkages/modifications on the array (Suppl Fig S4, S5). The data obtained with biotinylated BRs also agrees with previously published reports on GST-fusion BRs (19–22), and the comparable binding abilities of these biotinylated probes with the original BRs confirmed that biotinylation did not affect the Sia specificity of these proteins.

Considering the results with biotinylated-HsaBR on the sialoglycan microarray, we asked if HsaBR could be a replacement for MAL I and MAH, conventional lectins for this class of probes (SGRP3). In an experiment to compare Siaα2-3-binding preferences of Biotin-HsaBR, Biotin-MAL and Biotin-MAH, HsaBR showed significantly pronounced Siaα2-3-binding ability regardless of Sia modifications (*O*-acetylation, *O*-sulfation), or glycan structures (disaccharides, trisaccharides, oligosaccharides) (Suppl Fig S6A). Taken together with the glycan array data, we chose biotinylated HsaBR as our SGRP 3 probe (SGRP3*^Hsa^*)and and its non-binding variant ^Hsa^BR (R340E) as the nonbinding control (SGRP3*^Hsa^*NB) to characterize α2-3Sia-binding through various assays such as ELISA, Western Blotting, FACS and histochemistry, discussed below.

### SGRP4*^MHV^* as a probe for 4-OAc-Sias

A major contributor to mammalian sialoglycan diversity is *O*-acetylation, substituting the sialic acid hydroxyl groups at C4, C7, C8, and/or C9 (23, 24). The presence or absence of these *O*-acetyl moieties can block or promote binding of cellular and microbial lectins, and their regulation through sialate-O-acetylesterases (SOAEs) and sialate-O-transferases (SOATs) act as the molecular switches to control several cellular functions and interactions (25, 26). In contrast to *O*-acetylation at C7 or C9-OH, the occurrence of 4-*O*-acetylated Sia and its structural and functional significance have yet to be explored in much detail. In general, 4-OAc-Sias have been difficult to study even by chemical methods, due to variable expression or absence in many animal species, dynamic occurrence in pathological conditions, resistance to conventional sialidases, lability to acidic conditions (27) and masking of Sias from detection by some lectins. Equine erythrocytes, α2-macroglobulins and sera were used to study 4-OAc-“HD3”-reactive antibodies due to their high 4-*O*-acetyl-*N*-acetylneuraminic acid (Neu4,5Ac2) content (30– 50% of total Sias) (28). Guinea pigs serve as another common source of mammalian 4-OAc-Sias. Neu4,5Ac2 comprises a considerable share in serum (30% of all Sias) and liver (10% of all Sias) Sias in guinea pig, besides traces of 4-*O*-acetyl-*N*-glycolylneuraminic acid (Neu5Gc4Ac) in serum (29).

In humans, 4-OAc has been reported in tumor-associated antigens of colon cancers, melanomas and gastric cancers using “HD” antigen specific antibodies, for example a chicken antibody specific to Neu5Gc4Ac-lactosylceramide (4-OAc-HD3) recognized Neu5Gc4Ac in GM3 ganglioside fractions of human colon cancer tissues (30). Similar 4-OAc-HD3-reactive HD antibodies have also been reported in sera of patients suffering from malignancies and liver diseases (31). Two chicken MAbs HU/Ch2-7 and HU/Ch6-1 reacted with Neu5Gc4Ac in HD3 antigens (32). These MAbs, specific for individual glycan structure were a major tool for detection of 4-OAc in gangliosides or glycoproteins. Despite frequent reports on heterogenous 4-OAc in HD antigens in human cancers, there has not been a reliable conventional probe for *in situ* detection of this entire class of sialoglycans.

Previously, a sialic acid-binding lectin with specificity for *O*-acetyl Sia was purified from the hemolymph of the California coastal crab *Cancer antennarius*, which was more precise than other known lectins from horseshoe crab (*Limulus Polyphemus*) and slug (*Limax flavus*) but it also showed affinity towards 9-*O*-acetyl in addition to 4-*O*-acetyl Sias (33). A lectin from *Tritrichomonas foetus*, a parasitic protozoan that causes abortion in cows, was reported to react preferentially with Neu4,5Ac2 over de-*O*-acetylated Sias, and agglutinated equine erythrocytes containing Neu4,5Ac2 efficiently, but its preferences were also not exclusive for 4-O acetylated Sias (34).

In general, *O*-acetylation of Sias can be a major receptor determinant for some viruses. Among the first viruses shown to initiate infections via O-Ac-Sias were the influenza C viruses, human coronavirus OC43, Bovine Coronaviruses BCoV and porcine encephalomyelitis virus (PToV), but none of them exhibited binding of Neu4,5Ac2 (35–39). Infectious salmon anemia virus (ISAV), the causative agent of infections in Atlantic salmons showed specificity for, and hydrolysis of 4-OAcetylatedAc-Sias (40). ISAV preferentially de-*O*-acetylated free and glycosidically bound Neu4,5Ac2 and showed lower and no hydrolysis for free and bound Neu5,9Ac2 respectively. ISAV exhibited hydrolysis for both Neu4,5Ac2 and Neu4Ac5Gc at comparable efficacy, which was a significant advantage over known 4-OAc-Sia-binding molecules but it’s affinity for free Neu5,9Ac2 although lower than influenza C virus (40), restricted possibilities to derive a comprehensive probe for 4-OOAc-Sias.

Murine coronavirus mouse hepatitis virus (MHV-stain S) expresses a hemagglutinin-esterase that exhibits comparable sialate-4-O-Acetylesterase enzymatic activity to that of ISAV (41). Unlike comparable esterases from other sources, MHV-S HE protein specifically de-*O*-acetylates Neu4,5Ac2 but not Neu5,9Ac2, and converts glycosidically-bound Neu4,5Ac2-rich glycoproteins from horse and guinea pigs to Neu5Ac (41). MHV-S was able to hydrolyze acetyl esters from free as well as glycosidically-linked Neu4,5Ac2. Interestingly, MHV-A59 and several other MHV strains do not express a HE (42–44). Previously, the MHV-S HE ectodomain, released from HE-Fc by thrombin-cleavage, was reported to exhibit proper sialate-4-O-acetylesterase activity when assayed for substrate specificity with a synthetic di-*O*-acetylated Sia (4,9-di-*O*-acetyl-*N*-acetylneuraminic acid α-methylglycoside, αNeu4,5,9Ac_3_2Me) (45).

Considering MHV-S HE as a useful tool for the histological detection of Neu4,5Ac2, our collaborators had previously expressed the esterase-inactive MHV-S HE ectodomain as a fusion protein with a C-terminal Fc domain of human IgG1 and investigated Neu4,5Ac2 distribution in human and mouse tissues (46). In another study, we further modified the virolectin by fusing MHV-S-HE ectodomain-Fc to a hexahistidine (His6) sequence and detected high expressions of 4-OAc-Sias in horse and guinea pig respiratory tract tissues than mouse where it was mostly localized in gastrointestinal tract. 4-OAc-Sias were also found in a small number of cells within the duck, dog, and ferret respiratory tissues screened but not so far in the tissues of humans or pigs (47).

To derive a stable 4-OAc-Sia binding probe, we expressed the MHV-S-HE esterase inactive ectodomain (S45A), and the nonbinding mutant MHV-S-HE (F212A), as fusion proteins with the C-terminal Fc domain of human IgG1, along with Avi-tag for permanent biotinylation (Suppl Fig S3). Sia-binding and nonbinding proteins MHV-S-HE protein probes were biotinylated using manufacturer’s protocol and tested for their specificity for 4-OAc-Sias on sialoglycan microarray. As anticipated, biotinylated Sia-binding MHV-S-HE (S45A) exhibited very specific recognition of towards 4-OAc-Sias while the nonbinding mutant MHS-S HE (F212A) did not show any binding with any sialylated or non-sialylated glycan on the microarray, confirming its suitability as a non-binding control (Figure 1; Suppl Fig S4). Biotinylated MHV-S-HE (S45A) binds exclusively to Neu4,5Ac2α3Galβ3GalNAcαR1, Neu4,5Ac2α3Galβ3GalNAcβR1, Neu4,5Ac2α3Galβ3GlcNAcαR1, Neu4,5Ac2α3Galβ3GlcNAcβR1, Neu4,5Ac2α3Galβ4GlcβR1, Neu4,5Ac2α3Galβ4GlcNAcβR1 and Neu4,5Ac2α3Galβ4GlcNAcβ3Galβ4GlcβR1 (Figure 1). Despite such high specificity and avidity for Neu4,5Ac2, MHV-S-HE (S45A) did not show any binding of 4-OAc-Neu5Gc-glycans on the array (Fig 1). Currently, the sialoglycan microarray does not include α2-6-linked 4-OAc-Sias (Neu5Ac or Neu5Gc), hence the binding of MHV-S-HE α2-6-linked 4-OAc-Sias is not discussed here.

Considering the rare occurrence of Neu5Gc4Ac and limited knowledge about 4-OAc-Sia-recognizing proteins, MHV-S HE-derived fusion proteins provide the most useful probes for in situ detection of 4-OAc-Sias, represented mostly by Neu4,5Ac2. Given its exclusive preference for 4-OAcetylated-Sias, the biotinylated MHV-S-HE esterase inactive ectodomain appears to be the best probe for this class of sialoglycans (SGRP4*^MHV^* and SGRP4*^MHV^*NB) and demonstrate its utility through ELISA, Western blotting, FACS and histochemistry in this article. We expect that SGRP4*^MHV^* will facilitate research on functional significance of 4-OAc-Sias, help finding more comprehensive 4-OAc-Sia binding proteins and will allow us to seek an improved version of SGRP4 with recognition of both Neu4,5Ac2 and Neu5Gc4Ac.

### SGRP5*^IgY^* recognizes all Neu5Gc-terminated glycans

Humans are genetically defective in synthesizing the common mammalian Sia-Neu5Gc but can metabolically incorporate small amounts of this Sia from dietary sources into glycoproteins and glycolipids of human tumors, fetuses and some normal tissues (48). Neu5Gc derived from dietary sources has been observed in breast, ovarian, prostate, colon and lung cancers. Thus, there is need for sensitive and specific detection of Neu5Gc in human tissues and biotherapeutic products. Previously a number of different monoclonal antibodies against Neu5Gc have been reported which recognized Neu5Gc only in the context of particular underlying sequences, but those generally lack the ability to detect Neu5Gc on other structurally related or unrelated glycans (49–55).

We could not find a microbial Sia-binding protein that is completely specific for Neu5Gc, as there is always some cross-reactivity with few Neu5Ac glycans (51–53). Previously, we had improved the specificity and reduced background cross reactivity of a chicken polyclonal anti-Neu5Gc-glycan IgY by modifying the affinity purification procedures (56). Since then, this chicken anti-Neu5Gc IgY has been the only available probe for comprehensive *in situ* Neu5Gc-glycan detection.

To explore alternative Neu5Gc-specific probe from microbial sources, we previously reported Neu5Gc-binding preferences of subtilase cytotoxin (SubAB), an AB5 toxin secreted by Shiga toxigenic *Escherichia coli* (STEC) (10). SubB, the B5 subunits of STEC AB5 toxin, exhibits strong preference for Neu5Gc-terminating glycans and serves as key structural basis of STEC’s specificity towards Neu5Gc. SubB showed 20-fold less binding to Neu5Ac; over 30-fold less if the Neu5Gc linkage was changed from α2–3 to α2–6. Neu5Gc-binding by SubB suggested that a more selective diagnostic tool for Neu5Gc detection can be derived from the native protein. Day and colleagues, using molecular modeling and site directed mutations, managed to reduce the α2– 3 to α2–6-linkage preference, while maintaining or enhancing the selectivity of SubB for Neu5Gc over Neu5Ac (57). This SubB analog, SubB2M (SubBΔS106/ΔT107 mutant) did display improved specificity towards Neu5Gc, bound to α2–6-linked-Neu5Gc and could discriminate Neu5Gc-over Neu5Ac-glycoconjugates in glycan microarrays, surface plasmon resonance and ELISA assays. SubB2M also showed promising diagnostic properties in detecting Neu5Gc-glycans in early stages of ovarian cancer, and in late-stage diseases which express high levels of Neu5Gc-glycans (58).

Considering the promising Neu5Gc-binding properties of SubB proteins, we attempted to derive a comprehensive probe for Neu5Gc-glycans from SubB and tested His6-tagged versions of the reported SubB proteins including SubB, SubB2M and SubB2M-nonbinding mutant (SubB2M, S12A) on the glycan array. SubB2M exhibited significantly improved recognition of Neu5Gc-glycans compared to the native SubB, while the nonbinding mutant of Sub2M did not show any binding with glycans on the array (Suppl Fig S7). While we were able to reproduce the higher Neu5Gc preferences of Sub2M over native SubB, (59). our sialoglycan microarray data suggested that despite having promising Neu5Gc-binding properties, the Sub2M lacked the scope and robustness of Neu5Gc binding abilities of chicken anti-Neu5Gc IgY (Suppl Fig S7).

Considering results obtained from the Neu5Gc-binding molecules discussed or tested above, it can be inferred that there is no contemporary molecule better than the affinity-purified chicken polyclonal anti-Neu5Gc-IgY to detect a broader range of Neu5Gc-glycans. Hence, we selected chicken polyclonal specific anti-Neu5Gc-IgY as SGRP5 and biotinylated this along with its nonbinding control IgY (from non-immunized chickens) as the final set of SGRP5*^IgY^* (Suppl Fig S4). The utility of chicken Neu5Gc-IgY as SGRP5*^IgY^* in general lab-used methods including ELISA, Western Blotting, FACS and histochemistry is discussed in an earlier article, and the results obtained here confirmed its Neu5Gc-specificity as previously reported (56, 60). In a long run, it is necessary to identify a monoclonal IgY that can detect all forms of Neu5Gc-sialoglycans.

### SGRP6*^SNA^* recognizes all α2-6-Sias

Unlike bacteria, some viruses that cause upper respiratory infections such as human influenza A, B viruses, and human coronavirus OC43 exhibit preferential affinity towards α2-6Sias, which are abundant in the upper airway epithelial brush border in humans (61, 62). Influenza viral haemagglutinins (HA) are the major glycoproteins that allow the recognition of cells in the upper respiratory tract or erythrocytes by binding to α2-6Sias, and this property makes them potential candidates for α2-6Sias-binding probes. We investigated a range of viral HAs in the form of HA-Fc fusion proteins (soluble HA fused to human IgG1 Fc) for their sialoglycan-binding specificity using the glycan microarray (Suppl. Fig S8). Among the tested HA-Fc fusion proteins, Cali09 HA-Fc derived from California/04/2009 H1N1 showed selective binding to α2-6-linked Sias, most prominently with Neu5Acα6Galβ4GlcNAcβR5, followed by Neu5Acα6Galβ4GlcNAcβR1 and Neu5,9Ac2α6Galβ4GlcNAcβR1. Despite strong preferences for these α2-6Sias, none showed binding with a full range of α2-6Sias, specially α2-6Neu5Gc glycans. The failure of Cali09-HA-Fc to recognize a number of α2-6-linked sialosides, and its binding to a few α2-3 Sia-linked sialosides questioned its suitability as a probe for α2-6linked Sias (Suppl. Fig S8). Aichi68-HA-Fc derived from the hemagglutinin of Aichi/2/1968 H3N2 strain also lacked robustness and specificity showing indiscriminate binding towards a number of α2-6 and α2-3 Sias (Suppl. Fig S8). Surprisingly one candidate, PR8 HA-Fc derived from Influenza strain A/Puerto Rico/8/1934 (PR8 H1N1)), even showed prominent binding of α2-3-sialylated glycans instead of α2-6Sias (Suppl. Fig S8). Among other tested viral haemagglutinins, SC18 HA-Fc derived from influenza A H1N1 (A/SouthCarolina/1/18) exhibited selective binding to a few α2-6 Sias such as Neu5Acα6Galβ4GlcNAcβR5 and Neu5Acα6Galβ4GlcNAcβR1 while another virolectin Mem-HA-Fc from influenza A H1N1 (A/Memphis/1/1987) bound to α2-3Sias and α2-6Sias without any strong preference for either linkage (Suppl. Fig S8). Taken together, it can be inferred that the viral haemagglutinins tested exhibited promising specificity towards α2-6Sias but lacked the robustness and binding dynamics required for a probe. Our results and observations appear true for several other viral hemagglutinins not included in this study (63).

As an alternative α2-6Sia-specific microbial probe, we reviewed the α2-6Sia-binding properties of a recombinant lectin (PSL) from the mushroom *Polyporus squamosus* that was reported for its high affinity binding with Neu5Acα2-6Galβ4GlcNAc (64, 65). However, PSL showed a high order preference towards α2-6 over α2-3Sias but bound exclusively to N-linked recognizing Neu5Acα6GalNAc on serine/threonine-linked glycoproteins, we did not experimentally investigate this molecule further as an α2-6Sia-specific probe.

As an α 2-6Sia-binding alternative from mammals, we considered the well-characterized vertebrate sialic acid-dependent adhesion molecule CD22/Siglec-2 which is a member of the immunoglobulin superfamily that is expressed by B lymphocytes and binds specifically to Neu5Acα6Galβ4GlcNAc (66–70). hCD22-Fc (human CD22 fused with the Fc region of human IgG1) showed high affinity for a few α2-6Sias on microarray (Suppl Fig S9), but lacked the avidity required for a probe, particularly for α2-6-linked Neu5Gc. With a possibility to characterize 2 SGRP6 candidates; one specific for α2-6Neu5Ac and the other for α2-6Neu5Gc, we tried exploiting the evolutionary derived strong preference of mouse siglec-2 (mCD22) for α2-6Neu5Gc. mCD22-hFc showed exclusive binding towards α2-6Sias, but preferred Neu5Gc in general (Suppl Fig S9). Interestingly, 9-*O*-acetylation completely aborted hCD22’s binding to α2-6Neu5Ac (Suppl. Fig S9) but did not affect mCD22 binding to α2-6Sias. Nevertheless, the relatively poor avidity and inability of hCD22 to recognize O-Ac-substitution in α2-6Neu5Ac, and the non-specificity of mCD22 binding α2-3 sialyl-LNnT glycan (Neu5Gcα3Galβ4GlcNAcβ3Galβ4Glc) underlined their incompetency as an ideal α2-6Sia-binding probes, even in combination (Suppl Fig S9).

Among other molecules, we also analyzed the adhesin of non-typeable *Haemophilus influenzae* which was reported for high affinity towards α2-6-linked Neu5Ac (71). According to this study, HMW2 bound with high affinity to α2-6-linked Neu5Ac such as Neu5Acα6Galβ4GlcNAc (α2-6-sialyllactosamine) and could discriminate it from α2-3Sias (<120-fold lesser binding with α2-3 Siallylactosamine).-linked siallylactosamine). HMW2 indeed showed appreciable preference for the α2-6 over the α2-3-linkage but was not able to recognize α2-6Neu5Gc, disqualifying it from consideration as a comprehensive probe for α2-6Sias. In the absence of an evolutionarily derived microbial protein as a dynamic probe for α2-6Sias, we considered whether SNA (*Sambucus nigra* or elderberry bark lectin) is still the best available probe for α2-6Sias.

SNA exhibits high preference for the terminal Neu5Acα6Gal/GalNAc sequences in both N-linked and O-linked glycans (72). Due to its ability to discriminate Siaα6Gal/GalNAc from Siaα2-3Gal/GalNAc and ubiquitous binding to Neu5Ac/Gc with/without O-Ac substitutions, SNA has been the standard probe for studying α2-6Sias. Using Vector lab’s Biotin-SNA, we investigated SNA’s Sia-binding efficiency in sialoglycan microarray and observed very high preference towards α2-6Sias for both Neu5Ac and Neu5Gc and no measurable binding to α2-3Sias (Fig 1). Interestingly, the *O*-acetyl substitutions that were a major concern for hCD22, mCD22, viral hemagglutinins and HMW2, did not hinder SNA’s binding to α2-6Sias. Apart from its binding to 6-*O*-sulfated Galβ4GlcNAc (73), SNA displayed an exclusive preference for α2-6Sias in disaccharides, trisaccharides and oligosaccharides in Asn or Ser/Thr-linked glycans. There appears to be no better molecule than SNA to probe α2-6Sias, hence we selected Biotin-SNA as SGRP6 (SGRP6*^SNA^*) for this study. SGRP6*^SNA^* is the exception to the set of SGRPs in not having an internal control of specificity. We characterized SGRP6*^SNA^* as α2-6Sia-probe without any non-binding variant in routine lab assay methods in this article.

### SGRP7*^BCoV^* recognizes an exclusive class of 7,9-diOAc-Sias

Except for some claims of 7-*O*-mono-acetylated Sias in microorganisms (74, 75) and on human lymphocytes (76), 7-OAc-Sias are not commonly reported in natural glycans because of their instability. During synthesis, the primary attachment site of *O*-acetyl groups was exclusively to the hydroxyl at C-7 of Sias, and the ester group migrates from C-7 to C-9 (77). A hypothesis was proposed that SOAT enzyme would effectively be a 7-O-acetyltransferase incorporating O-acetyl groups primarily at C-7 of sialic acids, followed by their migration to the primary hydroxyl group at C-9 and further transfer of the acetyl residue to C-7, resulting in di- and tri-*O*-acetylated species (78). This suggests that in nature 7-OAc will be represented as 7,9-diOAc or 7,8,9-triOAc, but largely as 7,9-diOAc in mammalian sialoglycans. Considering, there is no stable 7-OAc, there may not be a true 7-OAc-Sia-binding protein, we defined specificities of SGRP7 to 7,9-diOAc-Sias. 7/9di,9-diOAc has traditionally been studied in bovine submaxillary mucin, where the amount of 7-OAc- and 7,9-diOAc-Sias combined is almost twice of 9-mono-OAc-Sias (46).

We also reviewed the sialoglycan specificity of lectins and microbial proteins, previously reported for their binding to 7,9-diOAc-Sias. Lectins from *Cancer antennarius, Achatina fulica* and *Tritichomonas foetus* showed higher preferences for 9-OAc and somewhat to 4-OAc but negligible to poor affinities for 7,9-diOAc-Sias (33, 34, 79). Viruses exhibit prominent binding to O-Ac-Sias but most show either 9-OAc preference (Influenza C viruses, human coronavirus OC43, porcine encephalomyelitis virus) or 4-OAc preference (Infectious salmon anemia virus, Puffinosis virus, mouse hepatitis virus strains-S, DVIM, JHM) binding over 7,9-diOAc (35–40, 46, 80, 81). The hemagglutinin esterases from bovine toroviruses (BTOV-B150, BToV-Breda), bovine coronaviruses (strains; Mebus and Lun) and equine coronavirus (ECoV-NC99) preferentially cleave 7,9-diOAc substrates over mono 4- or 9-OAc-substrates (46, 82). Bovine viruses exhibit selective binding towards 7,9-diOAc, particularly BCoV-Mebus having relatively pronounced preferences towards 7,9-diOAc variants of both Neu5Gc and Neu5Ac, but lower preferences for 9-OAcs (46). It is interesting that BCoV esterase selectively removes all 9-OAc residues in 7,9-diOAc-Sias in BSM, leaving 7-OAc attached. The residual 7-OAc residues do not attract binding of BCoV anymore, demonstrating that Sia 9-*O*-acetylation is a strict requirement and that mono 7-OAc-Sias do not serve as ligands (46). The S protein, another hemagglutinin in BCoV, showed high specificity exclusively for 9-OAc-Sias with no preference for 7-OAc and this raises possibility that HE is the only receptor-binding protein for 7,9-di-Ac in BCoV-Mebus (83, 84).

Since BCoV-Mebus-HE is the best characterized for 7,9-diOAc-Sia specificities, we expressed its esterase inactive mutant (BCoV-Mebus-HE S40A) and corresponding non-binding mutant (BCoV-Mebus-HE F211A), each fused to human IgG-Fc followed by a His6-tag and an Avi-tag. Both proteins were biotinylated with an Avi-tag and were tested for their sialoglycan preferences on sialoglycan microarray (Fig 1, Suppl Fig S4). As observed, the nonbinder mutant does not show any binding with any sialylated or non-sialylated glycans in the array while the Sia-binding molecule BCoV-Mebus-HE S40A exhibited some affinity for 9-OAc-Sias (Fig 1), agreeing with its Sia-binding pattern on sialoglycan microarray reported previously (46). However, similarly encouraging as the BCoV-Mebus-HE, S40A did not exhibit any binding to 4-OAc, most 9-OAc-Sias and non-acetylated sialyloligosaccharides, the sialoglycan microarray did not contain 7/9diOAc,9-diOAc-Sias, so it is not known whether the protein binds that form.

### SGRP8*^TeT^* binds to major class of (α2-8-linked) di-Sia oligosaccharides

Oligosaccharide sequences in gangliosides and a few glycoproteins that terminate in Sias, are not always mono-sialylated. In addition to α2-3 or α2-6-linkage with penultimate Gal, Sia can be linked with another Sia predominantly by α2-8-linkage forming α2-8-linked disialosides, α2-8 linked oligosialic and α2-8-linked polysialic acid chains. α2-8-linked disialyl structures are critical component of neuronal gangliosides and these disialyl gangliosides, specially GD3 and GD2 have been utilized as tumor markers for melanomas, gliomas and neuroblastomas (28, 30–32). Due to their utility in cancers either as biomarker for in vivo immunolocalization or in phase I and II trials to target disseminated neuroblastoma, a range of anti-disialoganglioside monoclonal antibodies have also been generated. For example, R24 is a mouse IgG3 monoclonal antibody (MAb) that reacts with the ganglioside GD3 expressed by cells of neuroectodermal origin (85). Similarly, a list of anti-GD2 specific MAbs; 3F8 (86), BW704 (87), 14G2a, 15G2a (88), AI-201, AI-287, AI-410. AI-425 (89) and anti-GD3 antibodies like MAbs AI-245 and AI-267 (89), Ch14.18 and Ch14.18/CHO have been characterized for their preferential binding to disialyl gangliosides. Further, MAbs JONES, D1.1 and 27A showed affinity for 9-OAc-GD3 but not 9-OAc-GD2 while Mab 8A2 binds both (90). Despite the large number of GD3 and GD2 MAbs, there has not been a single MAb that could serve as a comprehensive probe for the broader range of disialylated glycans.

Among mammalian proteins, myelin-associated glycoprotein (MAG) shows high binding for specific gangliosides in order of GQ1bα > GT1aα, GD1α > GD1a, GT1b >> GM3, GM4, but not for GM1, GD1b, GD3, and GQ1b. Despite high affinity for disia and oligosias, MAG’s binding to gangliosides is not exclusive for di-Sia linkages (91, 92). Siglec-7 is also reported to react mainly with disialyl structures such as disialyl Lewis α, disialyl galactosyl globoside, and ganglioside GD3, although it also lacks the probe like binding dynamics for this class (93). Among lectins reported to bind to disialylated oligosaccharides, few are *Cancer antennarius* lectin, *Agrocybe cylindracea* lectin and WGA. *Cancer antennarius* lectin binds with GD3 (94), *Agrocybe cylindracea* lectin binds with GD1a and GD1b while not clearly differentiating other structures containing NeuAcα3Galβ3GalNAc (95). WGA however preferred GT1 and GD1 over other gangliosides, exhibits very high affinity for monosialylated glycans and GlcNAc also, hence cannot be used as a di-Sia linkage specific probe (96).

Infectious microbes possess promising affinity for disialyl oligosaccharides in accordance with the glycan environment of their anchorage surfaces. Porcine sapelovirus (PSV) binds specifically to GD1α (97) and *Helicobacter pylori* recognizes GD2, GD1α and GD1β by its sialic acid-binding adhesin (SabA) (98). In agreement with high affinity for single ganglioside as observed in Vibrio cholerae (GM1), the majority of gut infecting bacteria and viruses exhibit binding with a specific sialoglycan structure, and their affinity cannot be generalized for range of disialyloligosaccharide sequences.

Tetanus (TeT) and botulinum neurotoxins (BoT) are produced by anaerobic bacteria *Clostridium tetani* and *Clostridium botulinum*, respectively, and share significant structural and functional similarity (99). Not only BoT (serotypes A to G) and TeT exhibit affinities for similar di and oligosialogangliosides but also share significant sequence homology. In both BoT and TeT, 150Kda single chain can be cleaved in to 50Kda N-terminal light chain and 100Kda C-terminal heavy chain. The HC fragment plays the primary role in receptor binding and can be further cleaved into 50Kda N-terminal fragment (HN) and 50Kda C-terminal fragment (HC). HC domain or HCR is characterized by the amino acid sequence homology among BoTs and TeT, which suggests that a conserved amino acid motif of in this domain may define a common carbohydrate recognition site. Since both BoT and TeT possess comparable binding specificity for gangliosides, we preferred TeT to investigate its potential for a comprehensive disialyl linkage recognizing probe. Selection of TeT over BoT was also based on detailed reports on TeT binding to gangliosides (100–103), and feasibility to select single serotype of TeT than BoT that has multiple serotypes. An optimal receptor-binding domain (residues 1105 -1315) from TeT-HCR was expressed as SNAPf-His6 fusion in the pGEX-3X vector in a bacterial expression system along with its mutants TeT-HCR R1226L, TeT-HCR W1289A, and TeT-HCR R1226L/W1289A (Suppl Fig S3 & S10). Expressed proteins were biotinylated by SNAP-Biotin chemistry as described in experimental procedures and tested for their sialoglycan affinity in microarray (Fig 1, Suppl Fig S4, S10). TeT has been shown to specifically bind gangliosides of the G1b series, GD1b or GT1b. In accordance with published report (102), TeT-HCR bound with a range of sialogangliosides including monosialylated GM1α, fucosyl-GM1 and oligosialylated GD1α, GD3, GT3 (Suppl Fig S10). The high affinity of TeT-HCR for GM1 and Fuc GM1 questioned its candidacy as di-Sia linkage specific probe. Single mutant; TeT-HCR R1226L and double mutant TeT-HCR R1226L/W1289A didn’t bind with any glycans on array, suggesting the significance of arginine residue at 1226 position in sialoglycan identification (Suppl Fig S4 and S10). Another single mutant TeT-HCR W1289A showed appreciable enhancement in affinity towards GD3, GT3 while loss in binding towards monosialylated and asialylated glycans on array (Suppl Fig S10). With respect to TeT-HCR (native protein), the tryptophan mutant TeT-HCR W1289A had pronounced specificity towards disialogangliosides structures particularly Neu5Acα8Neu5Acα3Galβ4GlcβR1, Neu5Acα8Neu5Acα3Galβ4GlcβR4, Neu5Acα8Neu5Acα8Neu5Acα3Galβ4GlcβR4, and Neu5,9Ac2α8Neu5Acα3Galβ4GlcβR1 (Suppl Fig S10). However, we, we noticed that TeT-HCR W1289A exhibited some minor binding with monosialogangliosides on the array but based on its high affinity for the range of α2-8-linked disialyl oligosaccharides and lack of other available options (MAbs, plant lectins, siglecs, viral, bacterial proteins) to derive a more potent probe for this class, we selected biotinylated TeT-HCR W1289A as SGRP8 (SGRP8*^TeT^*). We selected the double mutant; Biotinylated TeT-HCR R1226L/W1289A (SGRP8*^TeT^*NB) as a nonbinding control of SGRP8*^TeT^* due to its complete loss of binding for any glycans on array. The broad specificity of several bacterial toxin including TeT-HCR towards GM1, GD3, GT3 and GQ series of glycans could be related with evolutionary preference to bind with most available ligands in their neural niche. This nonexclusive binding with multiple ligands by toxins might have advantage for the microbes but reduces our possibility to derive a molecule with probe like robustness and dynamic specificity. Here, we present that site-specific modifications could improve the binding range in otherwise ‘nonspecific’ probe candidates and characterize SGRP8*^TeT^* as α2-8-disiaoligosaccharide binding probe through general laboratory methods later in this report.

### SGRP9*^PToV^* is a competent probe for 9-OAc-Sias

Of the variety of O-Ac-Sia modifications in nature, 9-*O*-acetylation is the most common in cells and tissues of humans and other animals. Their distribution is highly variable and implicated in embryonic development, host pathogen interactions and immunity. Although there have been monoclonal antibodies against *O*-acetylated gangliosides, these MAbs remained highly specific in recognition of *O*-acetyl esters only when represented by specific underlying sugar chain (104). Few Sia specific lectins recognizing 9-OAc modifications have been reported, for example a lectin from the marine crab *Cancer antennarius* showed affinity for 9-OAc and was used to demonstrate the presence of tumor-associated 9-OAc-GD3 on human melanoma cells (33). *Cancer antennarius* lectin also showed significant affinity for Neu4,5Ac2, and didn’t show affinity for 9-OAc. AchatininH, a lectin from the hemolymph of African land snail *Achatina fulica* did not bind to Neu4,5Ac2, but failed to exhibit probe-like preference for a number of 9-OAc-Sias (105). Another lectin from protozoan, *Tritichomonas foetus* showed promising affinities for 9-OAc-Sias but also bound to de-*O*-acetylated Sias relatively high affinity, confirming its unsuitability as a probe (34).

9-OAc-Sia-binding specific influenza C virus hemagglutinin esterase (Inf-CHE) was previously utilized as whole virions or recombinant protein for assessing a wide spectrum of sialoglycoconjugates like mucins, serum glycoproteins or gangliosides containing naturally or synthetically *O*-acetylated sialic acids (106). The influenza C virus hemagglutinin-esterase is a membrane-bound glycoprotein that binds specifically to 9-*O*-acetylated sialic acids (hemagglutinin activity) and then hydrolyzes the *O*-acetyl group (receptor-destroying activity). Inf-CHE can specifically cleave O-acetyl groups from Neu5,9Ac2 but not from 7-*O*-acetyl-*N*-acetylneuraminic acid (Neu5,7Ac2), and very slowly from Neu4,5Ac2 (107). Previously, we demonstrated that inactivation of Inf-CHE esterase by treatment of serine esterase inhibitor di-isopropyl fluorophosphate (DFP) resulted into a stable and irreversible binding with 9-OAc-Sias (106). However, using the whole virion as a probe was complicated due to variations in purity and stability of preparations, poor reproducibility, high nonspecific background, and lack of linearity in response. To avoid these limitations, we had replaced Inf-C virions with a soluble and recombinant chimeric molecule composed of the extracellular domain of Inf-CHE fused to the Fc region of human IgG1 (CHE-Fc). DFP inactivation of CHE-Fc stabilized the hemagglutinin activity and yielded a probe CHE-FcD that was more specific for 9-OAc-Sias than the whole Inf-C virion (108, 109). CHE-FcD provided a better alternative but did not qualify as a standard probe for 9-OAc-Sias due to selective preference for Neu5,9Ac2 over Neu9Ac5Gc glycans, nonspecific binding to Neu5,7Ac2 and hazards related to use of DFP (Supp. Fig S11), (40).

In order to derive a better probe for 9-OAc-Sias, we reviewed other O-Ac-esterases from Inf-C like mammalian coronaviruses; bovine (bovine coronaviruses; BCoV strains-Mebus, LUN, Breda, B150), birds (puffinosis virus), human (human coronaviruses, strains-OC43, HKU1), equine (ECoV-NC99), murine (mouse hepatitis virus strains-S, DVIM, JHM) and porcine (porcine torovirus strains; PToV, strains-Markelo, P4), reported previously (46, 80, 81). Based on these reports, which also includes our previous studies on OAc-Sia specificity of nidovirus HEs, we selected Porcine Torovirus P4 strain (PToV) hemagglutinin esterase to investigate as a potential probe for this class. Following the approach that we used for CHE-Fc, we expressed PToV HE ectodomain in HEK293T cells as fusion proteins with a C-terminal Fc domain of human IgG1 (PToV-HE-Fc). Instead of using DFP, we used site-directed mutagenesis (S46A) to inactivate the esterase activity in PToV-HE-Fc fusion protein. PToV-HE-Fc showed high selectivity towards 9-OAc-Sias and demonstrated its applicability to revealing the OAc-Sia distribution in human and animal tissues and cell lines (46, 47).

We expressed 9-OAc binding protein (PToV-HE-Fc, S46A) and nonbinding protein (PToV-HE-Fc, F271A) fused to human IgG-Fc followed by a His6 tag and an Avi-tag (Suppl. Fig S3). Both binding and nonbinding proteins were biotinylated using the Avi-tag and investigated for their affinities in a sialoglycan microarray. The esterase-inactive probe; PToV-HE-Fc (S46A) bound exclusively with 9-OAc-Sias including Neu5Ac and Neu5Gc, while hemagglutinin inactive nonbinding PToV-HE-Fc (F271A) did not show affinity towards any sialylated or nonsialylated glycans on array (Fig 1, Suppl Fig S4). PToV-HE-Fc (S46A) showed binding towards Neu5Ac and Neu5Gc α2-3, α2-6 and α2-8-linked to their penultimate sugars. In a similar sialoglycan microarray, CHE-FcD showed efficient binding with a number of Neu5,9Ac2-glycans but did not show similar affinity for Neu5Gc9Ac and remained a relatively weaker binder to 9-OAc-Sias in general (Suppl Fig S11). Alternatively, PToV-HE-Fc (S46A) exhibited significantly stronger binding towards a wide range of 9-OAc-Sia-glycans in comparison with CHE-FcD. Taking together, the efficacy, exclusivity and reproducibility of PToV-HE-Fc (S46A) as 9-OAc-Sia-binding protein, we selected biotinylated PToV-HE-Fc (S46A) as comprehensive probe (SGRP9*^PToV^*) for 9-OAc-Sia and biotinylated nonbinding PToV-HE-Fc (F271A) as SGRP9*^PToV^*NB, the internal control of SGRP9*^PToV^*’s specificity. We characterize the utility of PToV-HE-Fc (S46A) as SGRP9*^PToV^* in general laboratory methods in sections later in this article.

### Use of the optimized panel of SGRPs in common methods

Although the detailed sialoglycan microarray analysis provides insights into the Sia specificity of SGRPs, it is essential for SGRPs to exhibit equally precise affinities towards their ligands in ensemble of glycans represented in biological samples. In particular, the multi-antennary sialoglycans and structurally overlapping sialoglycans in biological samples may have a different interactions with probes than observed with individual sialoglycans in the glycan arrays. Qualitative Sia binding and ligand specificity of SGRPs were therefore tested by ELISA experiments with blood sera from 9 animal species, each signifying diverse sialoglycan composition in terms of presence or absence of N-glycolyl at C5 of Sia, *O*-acetylation, types of glycosidic bonds to penultimate sugars and overall oligosaccharide sequence. For example, mouse serum is proportionately high in Neu5Gc-Sias in comparison with *Cmah*^−/−^ mouse that remain exclusive for Neu5Ac-Sias and relatively higher in *O*-acetylation. Horse and guinea pig sera were included for their high representation of 4-OAc-Sias which was absent in other sera, as observed in HPLC analysis of Sias (Suppl. Table-1). Similarly, we included erythrocytes from similar 9 animal species, expecting to derive a relatable pattern of SGRPs binding with sera and RBCs. HPLC analysis of surface sialome of erythrocytes (Suppl. Table-2) suggests high variance in Sia diversity among animals which would be interesting to detect with SGRPs. We emphasized on minimal loss of heat and pH labile O-Acs during assays and modified the procedures accordingly.

### SGRP1*^YenB^* exhibits Sia specific binding to all mammalian sera

As results summarized in Fig 2 suggest, SGRP1*^YenB^* exhibits comparable affinity to all sera, indicating its broad range of Sia specificities. In accordance with the observation from glycan array (Fig. 1), SGRP1*^YenB^* exhibits strong binding towards sera whether it was Neu5Gc rich (mouse, goat) or Neu5Ac abundant (human, rat and *Cmah*^−/−^ mouse). Further, high proportions of Sia modifications such as 9-OAc substitutions (rat, rabbit and *Cmah*^−/−^ mouse) and more prominently, 4-OAc substitutions (guinea pig and horse serum) didn’t inhibit SGRP1*^YenB^*’s binding to serum sialoglycans (Fig. 2A). Sialoglycan microarray (Fig. 1) confirms SGRP1*^YenB^*’s strong binding towards sialoglycans that may directly be responsible for its binding to immobilized sera, but to exclude the potential contribution of serum protein interactions with SGRP1*^YenB^* from our observations, we performed binding assays with sera after partial and complete Sia depletion. Mild periodate oxidation of sera followed by borohydride reduction generates side-chain truncated-Sias with a terminal hydroxyl at 7^th^ carbon. SGRP1*^YenB^* showed significantly reduced binding to periodate oxidized sera that was proportional to the loss of desired Sia ligands after the truncation (Supp. Fig S12A). A question remained as to whether the residual binding of SGRP1*^YenB^* to sera, observed even after periodate treatments were due to SGRP1*^YenB^* interaction with non-sialylated structures in serum, or the *O*-acetyl substitutions protecting Sias from complete oxidation. When *O*-acetyl esters were removed by mild base treatment before periodate oxidation, the residual SGRP1*^YenB^* binding was completely abolished, thereby excluding the role of protein-protein interaction in SGRP1*^YenB^*’s binding to mammalian sera (Supp. Fig S12A). In the same experiment, base treatment without periodate oxidation did not significantly influence SGRP1*^YenB^* binding to sera, confirming the probe’s unbiased preference for Sias as seen in glycan array also (Fig 1). Significantly, the nonbinding variant of the probe, SGRP1*^YenB^* did not show any binding with any serum at all tested concentrations, as expected (Fig 2A).

**Figure 2.**
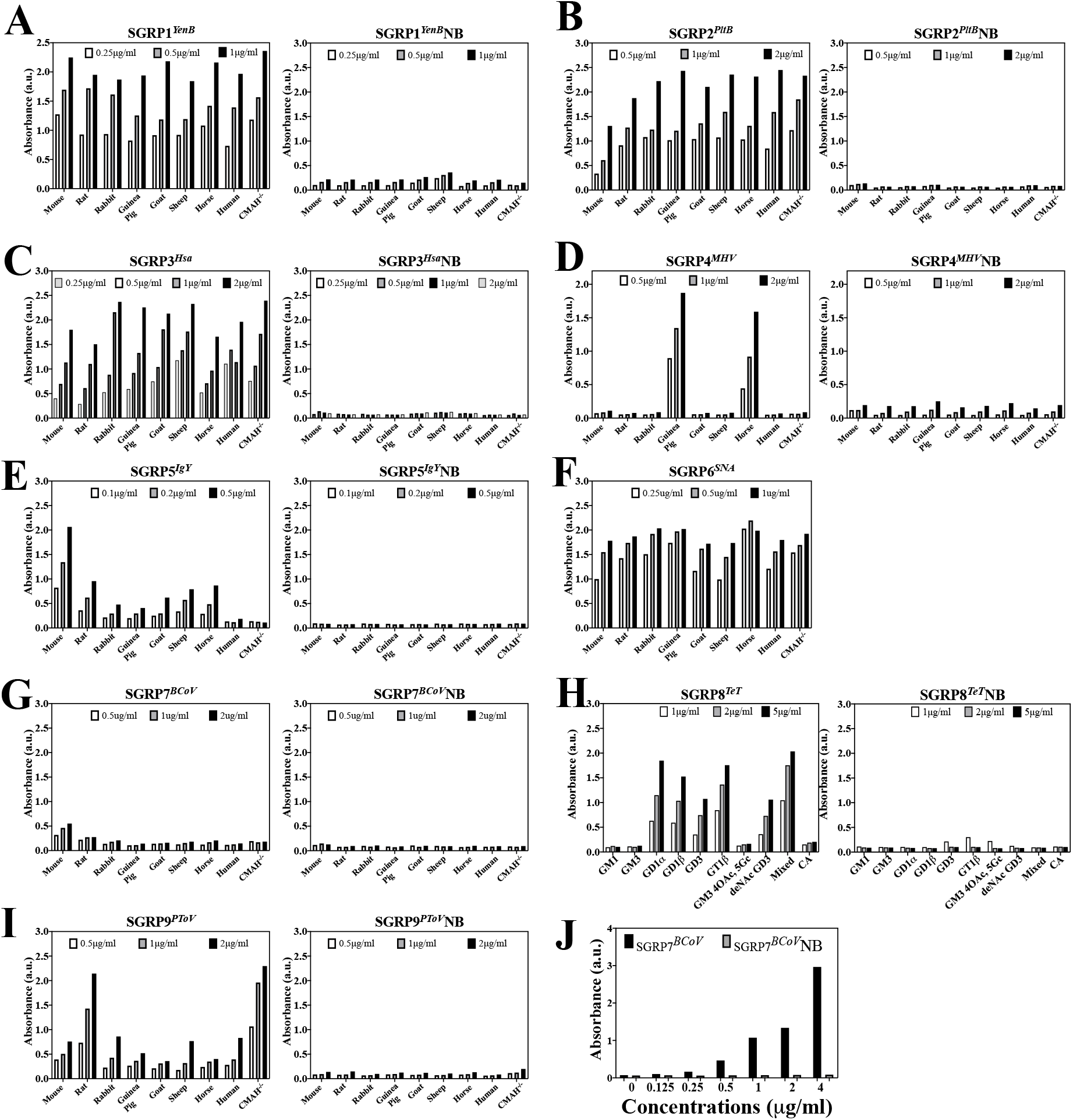
Specific binding of SGRPs to sialoglycans in mammalian sera. The figure presents SGRPs binding of sialoglycans in biological samples, here represented by mammalian sera from mouse, rat, rabbit, guinea pig, goat, sheep, horse, human and *Cmah*^−/−^ mouse. Panels A-I (except panel H) show binding efficiency of SGRPs and their nonbinding controls towards sera in concentration-dependent manner, as marked in individual panels. Panel H shows binding of SGRP8TeT and SGRP8TeTNB of purified gangliosides at concentrations mentioned in panel. Panel J demonstrates the 7,9-diOAc-Sia-specific binding of SGRP7BCoV with bovine submaxillary mucin while its mutant SGRP7BCoVNB shows no affinity towards BSM. Binding of biotinylated SGRPs was detected and developed using Avidin-HRP (1:1500). ELISA for each SGRP pair of was performed in same experiment/conditions and validity of binding was confirmed by background signals wells treated only with secondary antibody (Avidin-HRP), not shown here.

### SGRP2*^PltB^* exhibits Neu5Ac-specific binding to mammalian sera

SGRP2*^PltB^* binds to Neu5Ac and its derivatives that constitute a major fraction of Sia population in mammalian sera. Accordingly, SGRP2*^PltB^* exhibited binding to all serum types tested in a concentration dependent manner (Fig 2B). However, SGRP2*^PltB^* exhibits an interesting pattern of binding towards Neu5Gc rich mouse serum vs Neu5Ac rich sera, especially human and *Cmah*^−/−^ mouse sera. The non-binding mutant SGRP2*^PltB^*NB failed to bind serum from any animal source. Binding assays with mild base and periodate-treated sera confirmed that SGRP2*^PltB^*’s binding towards serum was exclusively to Sia and was devoid of nonspecific interactions (Suppl Fig. S12B). The SGRP2*^PltB^* also exhibited residual binding after periodate oxidation that were further reduced with prior base treatments, but base treatment alone did not noticeably influence the SGRP2*^PltB^* binding to sera.

### SGRP3*^Hsa^* exhibits α2-3Sia-specific binding to mammalian sera

SGRP3*^Hsa^* is specific towards Sias linked to underlying glycans with α2-3-linkage that coexist with α2-6Sias and α2-8-linked Sias in cells, tissues, and other biological samples. Among the sera employed to investigate SGRP3*^Hsa^* binding towards α2-3-linked Sias in complex biological samples, SGRP3*^Hsa^* bound all sera in a concentration dependent manner. In a different experiment to test Sia-specific binding of SGRP3*^Hsa^* to serum, the probe exhibited sera binding directly proportionate to their Sia constituents (Suppl Fig S12C), excluding non-specific interaction with nonsialylated structures in serum. Collectively, the results demonstrate SGRP3*^Hsa^*’s indiscriminate binding to α2-3 Sia in serum, irrespective of *O*-acetylation, Neu5Ac or Neu5Gc, a finding that agrees with our observations from sialoglycan microarray (Fig 1). SGRP3*^Hsa^*NB, the nonbinding version of SGRP3*^Hsa^* showed no binding with any serum at tested concentrations (Fig 2C).

### SGRP4*^MHV^* exhibits 4-OAc-Sia-specific binding to mammalian sera

SGRP4*^MHV^* represents a probe for an exclusive class of 4-*O*-acetylated sialoglycans, which show a high abundance in the blood components and tissues of certain animals. In our collection of sera, 4-OAc is represented by guinea pig and horse (38.7% and 32.4% of total Sia content respectively-Suppl. Table 1) and accordingly SGRP4*^MHV^* exhibited strong affinity towards these sera (Fig 2D). The negligible binding of SGRP4*^MHV^* to other sera suggests that its specific binding to guinea pig and horse serum was due it’s binding to 4-OAc-Sias exclusively found in these samples. Importantly, SGRP4*^MHV^* exhibited no binding to serum rich in 9-OAcs (*Cmah*^−/−^, rat and rabbit). confirming its ability to detect 4-OAcs over 9-OAcs. In similar experiment, SGRP4*^MHV^*NB didn’t bind to any serum at tested concentrations, confirming its utility as nonbinding control. In another qualitative approach to test SGRP4*^MHV^*’s specificity towards 4-OAcs, its binding was assessed with sera, pretreated with mild base and periodate. Periodate treatment did not have much effect on SGRP4*^MHV^*’s binding towards guinea pig and horse serum, whereas base treatment and base treatment followed by periodate oxidation completely blocked Sia detection (Suppl Fig S12D). These results, along with sialoglycan glycan microarray confirms that even in complex biological samples like serum, SGRP4*^MHV^* binds only to 4OAcSias

### SGRP5*^IgY^* exhibits Neu5Gc-specific binding to mammalian sera

Similar to SGRP4*^MHV^*, sera binding experiment for SGRP5*^IgY^* had internal indicators of probe’ specificity. While 5*N*-glycolylated Sias (Neu5Gc) are the major component of Sia diversity in mammals, Neu5Gc is absent from humans due to loss of functional *Cmah,* and also in *Cmah* ^−/−^ mice. The serum binding assay (Fig. 2E) exhibits that SGRP5*^IgY^* showed very strong affinity for the WT mouse serum, and also binding to all sera except human and *Cmah* ^−/−^ mouse. Significantly SGRP5*^IgY^* showed the reverse binding pattern to that was observed with SGRP2*^PltB^* (Fig 2B), suggesting an approach to double-check the specificity of such complimentary probes. SGRP5*^IgY^*NB didn’t bind any serum tested in similar experiment, as anticipated. A binding assay with base- and periodate-treated sera supported the specificity of SGRP5*^IgY^*, as removal of OAc esters did not affect its binding to serum, while removal of Sias abolished binding (Suppl Fig S12E).

### SGRP6*^SNA^* exhibits α2-6 specific binding to mammalian sera

SGRP6*^SNA^* exhibits strong and general binding with all mammalian sera in binding experiment, suggesting its robustness as α2-6Sia-recognition probe. As observed in glycan array (Fig1), SGRP6*^SNA^* does not discriminate between α2-6Neu5Ac and α2-6Neu5Gc, with or without OAc esters. Similarly, the serum binding data (Fig 2F) shows binding of SGRP6*^SNA^* to mouse, rabbit, guinea pig, human or *Cmah* ^−/−^ mouse sera which contain Neu5Gc, Neu5Ac & Neu5,9Ac2, Neu4,5Ac2, Neu5Ac, Neu5Ac and Neu5,9Ac2 respectively. SGRP6*^SNA^* has traditionally been used as α2-6 Sia recognizing probe, and our binding experiment with periodate oxidized serum confirms that SGRP6*^SNA^* binds to sera in Sia specific manner and does not bind serum proteins (Suppl Fig S12F).

### SGRP7*^BCoV^* exhibits 7,9-diOAc-Sia specific binding to Bovine Submaxillary Mucin

SGRP7*^BCoV^* exhibited insignificant binding to tested mammalian sera, likely due to very strong preference towards 7,9 diOAcSias, absent in these sera (Fig 2G). The probe does possess inconsistent and minor binding preferences towards 9-OAc-Sias but those were not detected in the sera sialoglycans. To confirm SGRP7*^BCoV^*’s recognition of 7, 9-diOAc-Sias, we performed binding experiments with immobilized BSM, enriched in di- and tri-OAc-sialoglycans. As anticipated, SGRP7*^BCoV^* showed dose-dependent binding patterns to BSM, while the nonbinding mutant SGRP7*^BCoV^*NB showed no detectable signals (Fig 2J). We confirmed its specificity towards OAcSiasOAc-Sias in base and mild periodate-oxidized BSM. The data (Suppl Fig S12G) explains that SGRP7*^BCoV^* binds BSM in OAc-dependent manner, and after depletion of OAc esters resulted in the dramatic reduction in SGRP7*^BCoV^*’s binding, comparable to loss of binding after complete depletion of Sias.

### SGRP8*^TeT^* exhibits DiSia linkage specific binding to gangliosides

SGRP8*^TeT^* exhibits very specific binding towards di-Sia linkages, the sialoglycan moieties mostly represented by gangliosides in biological system. As a result, probe SGRP8*^TeT^* showed no detectable binding with tested against sera in ELISA experiment (Suppl. Fig S13A). In an ELISA experiment with immobilized purified gangliosides, SGRP8*^TeT^* showed dose dependent binding towards di-sia oligosaccharide (GD1, GD3, GT1) discriminating from the mono-sia oligosaccharides (GM1, GM3) and polysia oligosaccharides conformations (Colominic acid) (Fig 2H). The observed results from ganglioside binding experiments are in complete agreement with the sialoglycan microarrays performed with SGRP8*^TeT^* and confirms the advantage of the probe over native TeT-HCR (Fig 1, Suppl. Fig S10 A & B). As anticipated, the nonbinding mutant SGRP8*^TeT^*NB showed no detectable signals, signifying its utility as control for SGRP8*^TeT^*’s specificity (Fig 2H).

### SGRP9*^PToV^* exhibits 9-OAc-Sia-specific binding to mammalian sera

SGRP9*^PToV^* binds to 9-OAc-Sias that are represented by all mammalian sera included for binding assays (Suppl Table-1). In observed results, SGRP9*^PToV^* demonstrated remarkably high affinity towards rat and *Cmah*^−/−^ mouse serum, suggesting their high proportions of Neu5,9Ac2 (Fig 2I). However, relatively lower binding to rabbit serum, as detected by signals in ELISA experiment was interesting as rabbit serum contained the highest fraction of Neu5,9Ac2 among the tested sera, but the basis of such variance is not known. It is clear from the serum-binding data of the SGRPs that those can demonstrate the presence or absence of the preferred ligand, but that this may not always measure the proportions relative to other Sia types within the serum. No detectable signals from similar binding assay with SGRP9*^PToV^*NB confirms that SGRP9*^PToV^* binding was specific. As observed in different ELISA experiments with OAc depleted sera, SGRP9*^PToV^*’s binding to mammalian sera including rat and *Cmah*^−/−^ mouse showed a linear and reproducible binding exclusive to base labile OAc esters (Suppl Fig S12H). The data showed that depletion of non-O-acetylated Sias doesn’t influence SGRP9*^PToV^* binding significantly while removal of OAc esters from Sias, or complete diminution of Sia population abolished its binding to sera (Suppl Fig S12H). These results together with sialoglycan microarray data (Fig1) confirm that SGRP9*^PToV^* exhibits very strong preferences towards 9OAcSias and does not interact with nonsialylated structures or proteins in complex biological samples.

### Western Blot analysis of SGRPs specificity towards sialoglycans in mammalian sera

To confirm that SGRPs are applicable to routine methods of glycoprotein analysis, we assessed SGRP’s qualitative detection of sialoglycoproteins by western blots. The gel electrophoresis and Western blotting protocol was modified to minimize loss of sensitive OAc-Sias (110, 111). As observed, SGRP1*^YenB^* bound to all 9 sera included in the experiment, which agrees with the observation from serum binding ELISA experiments (Fig. 3A). SGRP2*^PltB^* also bound to sera in accordance with their Neu5Ac contents, with a noticeable difference between its affinity for WT and *Cmah*^−/−^ mouse sera (Fig. 3C). SGRP3*^Hsa^* and SGRP6*^SNA^* showed binding to all sera, a pattern that was also observed in serum binding ELISAs (Fig. 3E & 3K). SGRP4*^MHV^* exclusively bound to guinea pig and horse sera (Fig. 3G), while SGRP9*^PToV^* exhibited high binding towards rat, rabbit and *Cmah*^−/−^ mouse sera (Fig. 3N), directly corresponding to OAc-Sia contents of these sera (Suppl Table 1). Interestingly, SGRP7*^BCoV^* showed significant binding with multiple sera which differs from our observations with sera binding ELISA experiments. In ELISA, SGRP7*^BCoV^* showed insignificant binding to sera and the overall binding pattern varied between experiments, suggesting a lack of linear binding behavior. We speculate that changes in pH and temperature during gel electrophoresis or blotting may result in migration of OAc esters, resulting in binding in western blotting, but we did not investigate this in detail. Blots, interrogated with non-binding variants of SGRPs (Fig. 3) did not show any binding to sera, suggesting Sia specific binding of SGRPs to sera. As SGRP8*^TeT^* did not bind sera in ELISA it was excluded from western blot analysis of serum proteins.

**Figure 3.**
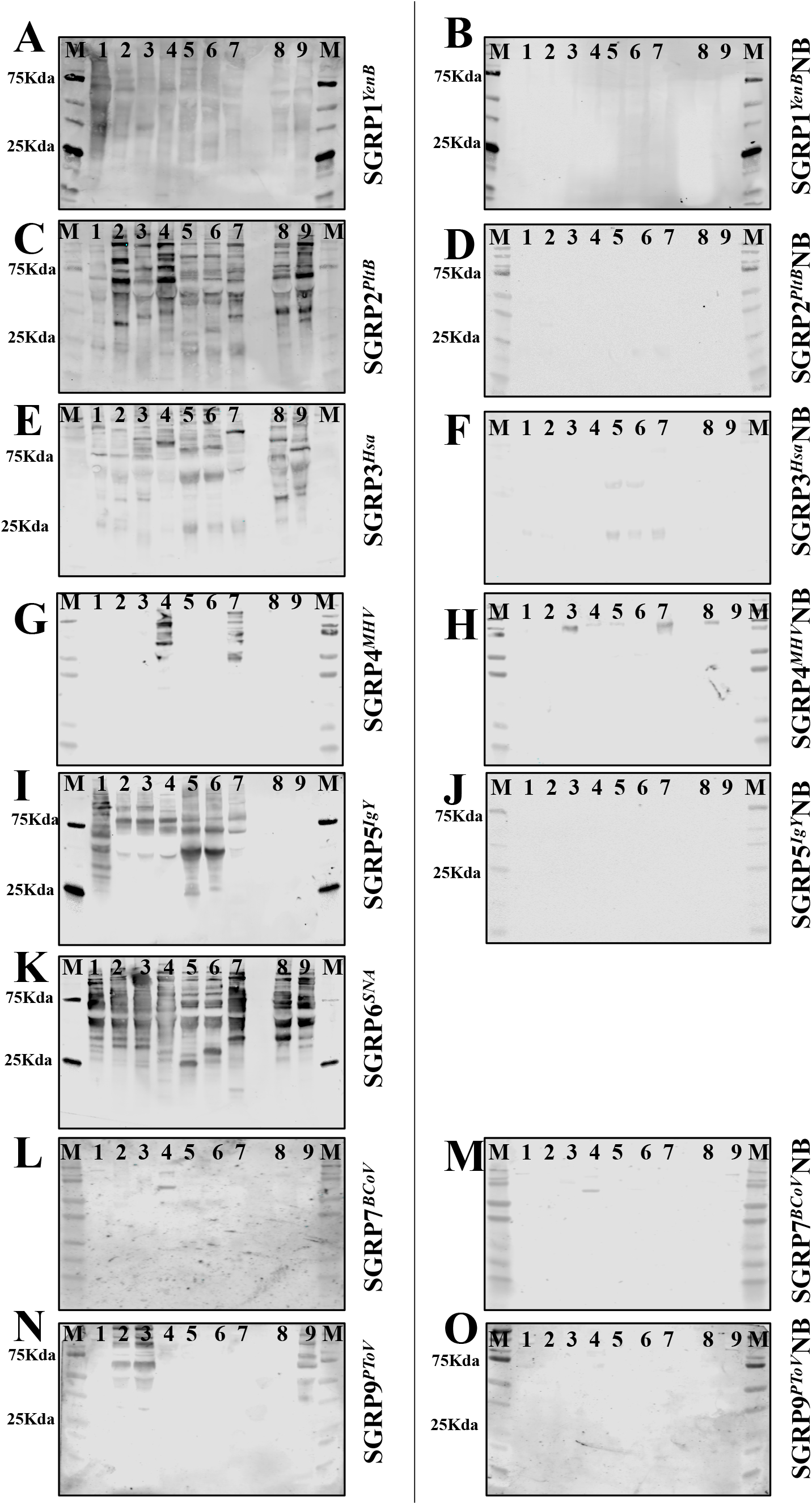
Specific recognition of sialoglycoproteins in mammalian sera by SGRPs in western blots. Mammalian sera from mouse (1), rat (2), rabbit (3), guinea pig (4), goat (5), sheep (6), horse (7), human (8) and *Cmah*^−/−^ mouse (9), were subjected to SDS-PAGE under conditions that protect labile O-acetyl groups immunoblotted and probed with SGRPs. M in each blot represents the molecular weight marker. Panels A & B show blots interrogated with SGRP1YenB and SGRP1YenBNB (both at 1 µg/ml); panels C & D show blots interrogated with 1 µg/ml SGRP2PltB and SGRP2PltBNB; panels E & F represent SGRP3Hsa and SGRP3HsaNB (both at 1 µg/ml); panels G & H shows blots probed with 1 µg/ml of SGRP4MHV and SGRP4MHVNB; panels I & J shows blot probed with 0.33 µg/ml SGRP5IgY and SGRP5IgYNB; panels K is blot interrogated by 1µg/ml SGRP6SNA; panels L & M show blots probed by 1 µg/ml SGRP7BCoV and SGRP7BCoVNB, and panels N & O represent blots interrogated with 2 µg/ml SGRP9PToV and SGRP9PToVNB. Binding of biotinylated SGRPs was detectable and developed using Streptavidin IRDye 680 (1:10,000). Analysis was done on a LiCor Odyssey infrared imager. Both blots for each SGRP set were detected at the same time using the same conditions.

### SGRP binding to frozen tissue sections: testing SGRP’s utility through *in situ* detection of Sias

The panel shows the typical binding patterns observed with each of the probes (red color indicates binding) (Fig. 4). The non-binding control shows no binding to any of the sections, as expected. SGRP1*^YenB^* detects mucins and blood vessels in many organs, and this is best demonstrated in the sections of kidney, where the glomeruli are highlighted with this probe (Fig 4A). The glomeruli, as well as the capillary blood vessels between the tubules are prominent in the wild type mouse, while in the *Cmah*^−/−^ mouse the capillaries between tubules are not as prominent (Fig 4B). The non-binding control shows no binding to any of the sections, as expected (Fig 4C). SGRP2*^PltB^* detects mucins and blood vessels in many organs, and this is best demonstrated in the sections of kidney, where the glomeruli are highlighted, and also the capillaries in between the tubules are visible (Fig 4D & E). The non-binding control shows no binding to any of the sections, as expected (Fig 4F). SGRP3*^Hsa^* detects mucins and blood vessels in many organs, and this is best demonstrated in the sections of kidney, where the glomeruli are prominently highlighted (Fig 4G). The binding to the kidney glomeruli in *Cmah*^−/−^ mouse is slightly more prominent (Fig 4H). The non-binding control shows no binding to any of the sections, as expected (Fig 4I). SGRP4*^MHV^* showed selective detection of mucins, and this is best demonstrated here in the mucin contained within the acini of pancreas. Staining was faint in the wild type and was much more prominently identified in the *Cmah* null animal (Fig 4J & K). The islets of the pancreas (the endocrine portion) and the non-binding control show no evidence of binding (Fig 4L). SGRP5*^IgY^* detects blood vessels in many organs and demonstrated here by the detection of the capillaries within the sections from the pancreas in wild type, but not in *Cmah*^−/−^mice (Fig 4M & N). The non-binding control shows no binding to any of the sections (Fig 4O). SGRP6*^SNA^* detects mucins and blood vessels in many organs. Here we observe that SGRP6*^SNA^* detects blood vessels in the pancreas of the *Cmah*^−/−^mouse much better than it does in the wild type animal (Fig 4P & Q). The tissue sections were incubated with secondary antibody Cy3-SA and served as control for SGRP6*^SNA^* specificity, and shows no binding to any of the sections, as expected (Fig 4R). SGRP7*^BCoV^* detects blood vessels in some organs but also detected white matter in the brain of the *Cmah*^−/−^mouse extremely well (Fig 4T). The non-binding control shows no binding to any of the sections. (Fig 4U). SGRP8*^TeT^* detected large areas of the non-nuclear neuropil in the brain from both the wild type and in the *Cmah*^−/−^ mouse. The white matter in the brain from the *Cmah*^−/−^mouse was even more prominent, and the non-binding control showed no binding to any of the sections (Fig 4V, W & X). SGRP9*^PToV^* detected mucins and also red blood cells in many of the organs However, in sections of liver from the *Cmah*^−/−^mouse, there was an abnormally high expression in the sinusoidal endothelial cells around the central vein, indicating right heart failure in the *Cmah*^−/−^mouse (the portal triads, have two sources of blood supply and are thus the better perfused areas in the liver) (Fig 4Y & Z). The non-binding control showed no binding to any of the sections, as expected (Fig 4Z1).

**Figure 4.**
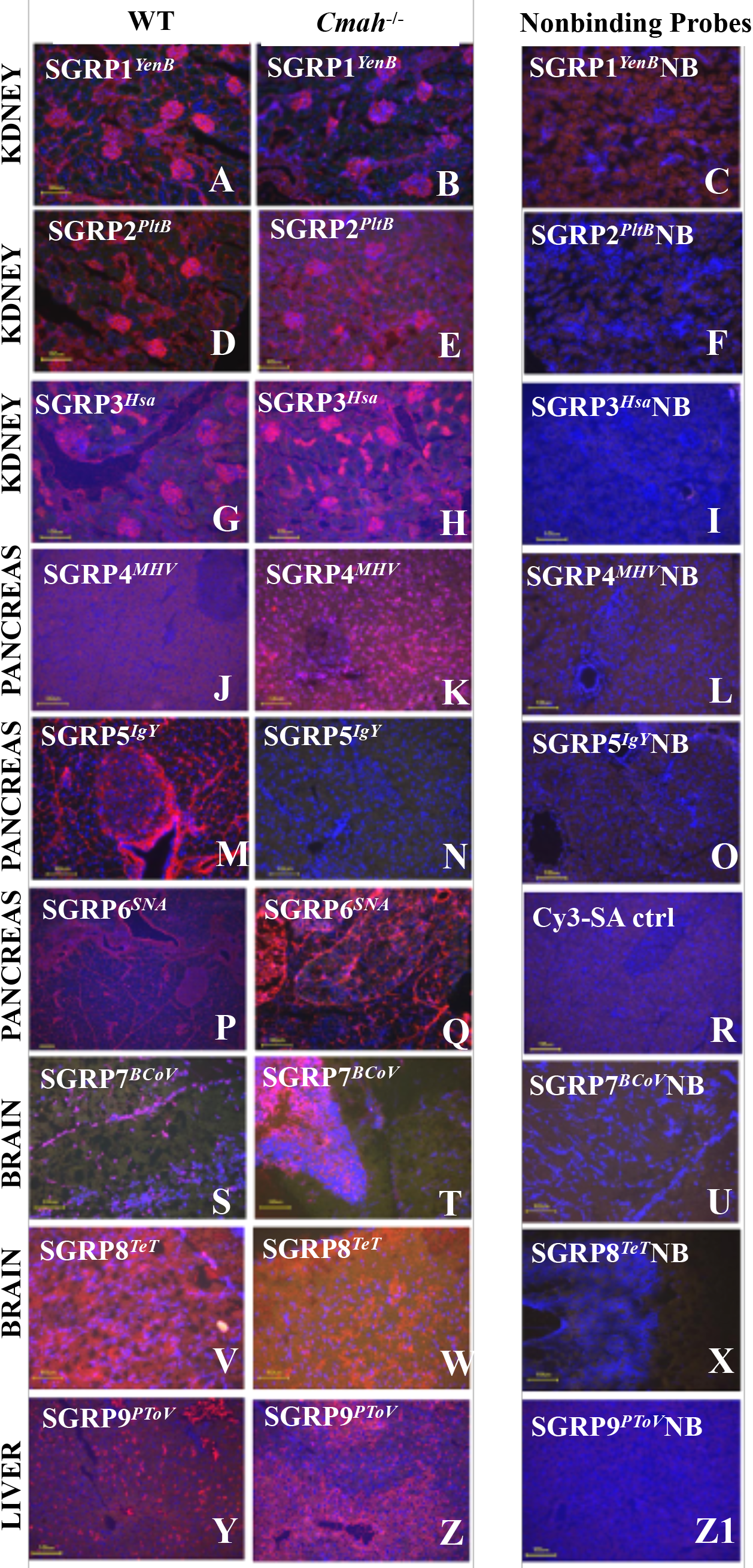
Histological analysis of frozen mouse tissues by SGRPs: The panel shows examples of binding patterns observed with each SGRP; binding indicated by red fluorescent staining. *In situ* sialoglycan detection ability of SGRP1*^YenB^* (5 μg/ml) is demonstrated using frozen kidney sections from WT and *Cmah*^−/−^ mouse (*Panels A & B*) and showing binding on glomeruli and blood vessels while the nonbinding variant SGRP1*^YenB^*NB shows no binding to tissue sections as expected (*Panel* C). *Panels D, E & G, H* show SGRP2*^PltB^* (5 μg/ml) and SGRP3*^Hsa^* (3 μg/ml) recognition of glomeruli and vessels in kidney sections from WT and *Cmah*^−/−^ mouse, while *Panel F & I* show no binding of SGRP2*^PltB^*NB and SGRP3*^Hsa^*NB to tissue sections as expected. SGRP4*^MHV^* (5 μg/ml) shows selective detection of its ligand in mucins in the pancreatic acini more prominently in *Cmah*^−/−^ mouse (*Panel* K) than in WT (*Panel J*), while SGRP4*^MHV^*NB shows no evidence of binding at all as expected (*Panel L*). *Panel M* (1 μg/ml) shows SGRP5*^IgY^* binding to WT mouse pancreas, while *Panel N* shows no binding, with the expected lack of SGRP5*^IgY^* ligands in *Cmah^−/−^* mouse. SGRP5*^IgY^*NB does not bind to tissue sections (*Panel O*) as expected. *Panels P & Q* represent SGRP6*^SNA^* (2 μg/ml) binding to blood vessels in WT and *Cmah^−/−^* mouse pancreas. *Panel R* represents a control experiment; tissue stained with secondary antibody Cy3-SA (1:500) only. *Panel S & T* shows that SGRP7*^BCoV^* (10 μg/ml) binds to blood vessels in brains of WT and *Cmah^−/−^* mouse and *Panel U* demonstrate no binding of SGRP7*^BCoV^*NB in same tissue as expected. *Panel V & W* suggest that SGRP8*^TeT^* (5 μg/ml) binds to its ligand predominantly present in WT and *Cmah^−/−^* mouse brains, while SGRP8*^TeT^*NB doesn’t show any binding on tissue section (*Panel X*). *Panels Y & Z* represent SGRP9*^PToV^* (5 μg/ml) binding to livers of WT and more prominently in livers of *Cmah^−/−^* mouse. *Panel Z1* shows SGRP9*^PToV^*NB doesn’t binding to tissue, as expected.

### SGRP binding to mammalian erythrocytes in flow cytometry

We performed flow cytometry experiments with mammalian RBCs to demonstrate *in situ* detection of extracellular Sias by SGRPs (Fig. 5). As shown in Fig 5A, SGRP1*^YenB^* bound with RBCs from all 9 animal sources tested, while its nonbinding version SGRP1*^YenB^*NB didn’t show any binding (Fig 5A). In general, the total fluorescence intensity for nonbinder probes remained comparable to autofluorescence (RBCs without any treatment) and antibody control (RBCs detected with SA-PE only). SGRP2*^PltB^* bound with all RBCs except wild type mouse. The most robust bindings of SGRP2*^PltB^* were detected with *Cmah*^−/−^mouse, guinea pig, rabbit and rat followed by goat and human RBCs (Fig 5B). SGRP2*^PltB^*NB didn’t show any noticeable binding with RBCs from any source. SGRP3*^Hsa^* bound with all types of RBCs with maximum binding detected from guinea pig, goat and rat RBCs and minimum with rabbit, indicating variable abundance of α2-3Sias on these erythrocytes (Fig 5C). SGRP3*^Hsa^*NB didn’t bind to any RBC type significantly. SGRP4*^MHV^* showed no noticeable binding to any RBC included in experiment, and the binding as detected from SGRP4*^MHV^* and SGRP4*^MHV^*NB remained comparable, suggesting absence of 4OAcSias in extracellular surface of these RBCs (Fig 5D). SGRP5*^IgY^* bound with all RBCs except from *Cmah*^−/−^ mouse as anticipated (Fig 5E). Remarkably, SGRP5*^IgY^* also didn’t bind strongly to guinea pig and rat RBCs, suggesting less abundant Neu5Gc in these RBCs in comparison to mouse, horse and rabbit. SGRP5*^IgY^*NB showed negligible detection, as expected from a nonbinding control probe. A stronger SGRP6*^SNA^* binding was detected with human, guinea pig and rabbit in comparison to goat, sheep and horse (Fig 5F). Taking together with SGRP3*^Hsa^*’s binding to RBCs, the relative abundance of α2-3Sias over α2-6Sias can be inferred for certain mammalian RBCs like goat, sheep and guinea pig. In absence of nonbinding mutant of SGRP6*^SNA^*, the results are provided with autofluorescence and fluorescence antibody controls. The 7,9-diOAc-Sia-specific binder SGRP7*^BCoV^* showed no appreciable binding in flow cytometry experiment, suggesting undetectable amounts of its ligand in tested RBCs (Fig 5G). The resultant binding of SGRP7*^BCoV^* and SGRP7*^BCOV^*NB remains comparable with RBCs; the results showed agreement with probe’s response with serum ELISA experiment, suggesting lack of SGRP7*^BCoV^* binding in absence of its exclusive ligand 7,9-diOAc-Sia. α2-8-linked disialyl-oligosaccharides specific probe SGRP8*^TeT^* didn’t show any binding with RBCs tested (Suppl Fig S13B) and hence a set of GD3 transfected CHOK1 cells were used to investigate efficiency of SGRP8*^TeT^* in flow cytometry experiment. As shown in Fig 5I, SGRP8*^TeT^* exhibited appreciable binding to GD3 expressing CHOK1 cells (CHOK1-GD3^+/+^) with respect to normal CHOK1 cells. The nonbinding SGRP8*^TeT^*NB showed relatively minor binding with cells, that was not affected by expression of GD3 gangliosides. As presented in Fig 5H, SGRP9*^PToV^* showed remarkable binding with rat RBCs among all tested erythrocytes and its nonbinder probe SGRP9*^PToV^*NB remained completely undetectable throughout the experiment. This experiment demonstrates the utility of SGRPs in characterization of the extracellular sialome of cells -for example showing that rat RBCs possess a dense Sia population that contains varying amounts of in α2-3-Sias, Neu5Ac and 9OAcSias9-OAc-Sias than α2-6-Sias, Neu5Gc and 4OAc4-OAc-Sias.

**Figure 5.**
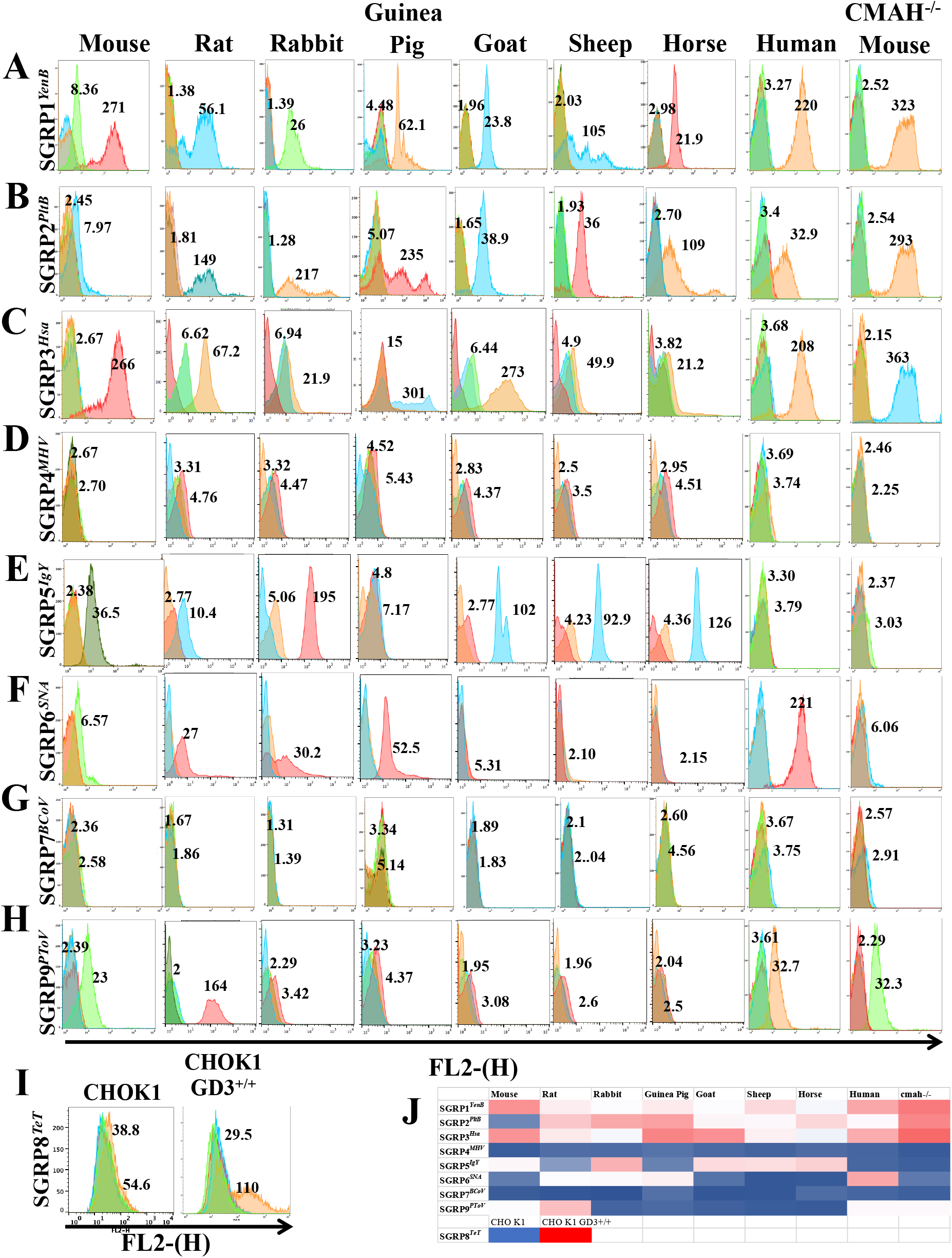
SGRPs recognize Specific sialoglycans on mammalian erythrocytes: The panels, as marked in figure represent SGRPs binding with RBCs for 9 animal sources. The mean fluorescence intensities (MFI) values for SGRPs; binding and non-binding versions are mentioned for each type of erythrocyte. Panel A represents SGRP1*^YenB^* (5 μg/ml) binding with all tested RBCs. Panel B represents SGRP2*^PltB^* (5 μg/ml) binding with RBCs in accordance to the fractional representation of Neu5Ac-sialoglycans. Panel C shows SGRP3*^Hsa^* (5 μg/ml) binding with all RBCs, while Panel D suggest SGRP4*^MHV^* does not specifically bind to RBCs, suggesting lack of its ligand. SGRP5*^IgY^* (2 μg/ml) binds prominently with all RBCs except rat and guinea pig (Panel E) while SGRP6*^SNA^* (2 μg/ml) lacks binding to goat, sheep and horse RBCs (Panel F). Panel G suggest the RBCs do not express SGRP7*^BCoV^*’s specific ligand 7,9-diOAc-Sias, resulting into negligible binding of the probe (5 μg/ml). Panel H shows SGRP9*^PToV^* (5 μg/ml) affinity towards surface glycans terminating in 9-OAc-Sias on RBC surfaces. Panel I represent significantly enhanced binding of SGRP8TeT (5 μg/ml) towards disialogangliosides expressed by GD3 transfected CHO cell in comparison to native CHOK1 cells. Panel J is the comprehensive heat map representation of SGRP’s binding to mammalian erythrocytes; color red and blue represent maximum and minimum binding efficiencies respectively.

## Conclusions and Perspectives

In this study we exploited naturally evolved Sia-recognizing microbial proteins to develop a feasible approach to design and derive sialoglycan-detecting probes, and applied the approach to develop and characterize a set of practicable probes recognizing the most predominant classes of mammalian Sia types. The study also reviewed and re-examined current standing of traditional probes or probe-like molecules that have been reported against known diversity of mammalian Sia types. We reviewed, assessed, and experimentally analyzed several probes to compare their suitability for use as updated SGRPs. In this process, we successfully identified an alternative for *Maackia* lectins in the form of SGRP3*^Hsa^*. The report does underline the utility of conventional α2-6Sia-binding *Sambucus nigra* lectin which appeared to be better than several microbial and mammalian proteins tested or reviewed in this study. Among developed probes; SGRP2*^PltB^*, SGRP4*^MHV^*, SGRP8*^TeT^* and SGRP9*^PToV^* represent novel probes for their class of Sia molecules, while SGRP1*^YenB^* and SGRP3*^Hsa^* showed substantial improvement over previously known probes. SGRP5*^IgY^* was also included in the final set, and SGRP6*^SNA^* remained unique as we could not identify better alternative from microbial sources. Neu5Gc-specific microbial proteins showed utility but could not match the specificity and robustness of anti-Neu5Gc IgY, that was finally selected as SGRP5*^IgY^*. An exception to SGRPs set, SGRP7*^BCoV^* is characterized as an imperfect probe that is not completely selective for 7OAcSias, and that must be updated with more precise SGRP7 if one is identified in the future. Similarly, if a new probe is developed that is specific for Neu5Gc, that would replace the SGRP5*^IgY^*which is currently selective for 7-OAc-Sias.

There are efforts ongoing by other groups, both in academic and industry to provide state of art sialoglycan detecting probes, and we attempted to review of most such studies and theoretically compared SGRPs with those molecules. Our focus remained on merits of proposed SGRPs and their utilities for a common biological science investigator. To have a consolidated assumption about the ‘best’ available probe for any given sialoglycan class, a real time comparison is required which is outside the scope of this study. In conclusion, this study provides a comprehensive set of probes for mammalian Sia types, designed and developed for the non-sialic acid experts. Several tests of demonstration of SGRPs specificity confirms their utility as practicable probes which can, and we expect SGRPs to be adapted by both experts and non-experts as tools of Sia recognition in their studies.

## Experimental Procedures

### Preparation of constructs

#### SGRP-1 – Y. enterocolitica Toxin B5; Y. enterocolitica Toxin B5 (Non-Binding)

The full-length coding region of YESubB (423 bp) incorporating a 3’ Hisx6 tag in the bacterial expression vector pBAD18 was kindly provided by James Paton and is described elsewhere. The corresponding non-binding mutant YESubB (S31A; Y100F) was engineered from the wt template above using the Q5 Site-Directed Mutagenesis Kit (New England BioLabs) and sequence verified.

#### SGRP-2 – S. typhi toxin B5; S. typhi toxin B5 (Non-Binding)

The Toxoid sequence pltB_pltA(E133A)_cdtB(H160Q)-6xHis and its corresponding non-binding control pltB(S35A)_pltA_cdtB-6xHis in the bacterial expression vector pET28b (Novagen) was kindly provided by Jorge Galan and described elsewhere. PltB or PltB(S35A) (411 bp) were amplified from the above respective constructs by PCR using pfu polymerase and primers 5’ GCGCG<u>CCATGG</u>ATGTATATGAGTAAGTATGTACC 3’; Nco1 site underlined and 5’ ATATA<u>AAGCTT</u>CTTGGGTCCAAAGCATTGTGTCG; Hindlll site underlined. PCR products were digested with Nco1 and Hindlll and sub-cloned into the corresponding sites of pET28b. Constructs were sequence verified.

#### SGRP-3 – S. gordonii hemagglutinin surface adhesion BR; S. gordonii hemagglutinin surface adhesion BR (Non-Binding)

The HsaBR sequence and its corresponding non-binding control HsaBR(R340E) (709 bp) in the bacterial expression vector pGEX-3X (Millipore Sigma) was cloned by Paul Sullam’s Laboratory and described elsewhere. HsaBR/pGEX-3X and HsaBR(R340E)/pGEX-3X were subsequently modified to incorporate a 3’ SNAPf_Hisx6 tag. Briefly, a Hisx6 sequence tag was inserted into the BamH1 and Xho1 sites of MCS2 of pSNAPf (New England Biolabs) to create pSNAPf_Hisx6. SNAPf_6xHis was amplified from this construct by PCR using the forward primer 5’-AAAA<u>GAATTC</u>AACCATGGACAAAGACTGCGAAA-3’ and reverse primer 5’-TTTT<u>GAATTC</u>ATTAGTGATGGTGATGGTGATGGGA-3’; EcoR1 site underlined. PCR products were digested with EcoR1 and sub-cloned into the EcoR1 site of HsaBR/pGEX-3X and HsaBR(R340E)/pGEX-3X above. Constructs were sequence verified.

#### SGRP-4 – Mouse Hepatitis Virus-S Hemagglutinin Esterase (Inactive); Mouse Hepatitis Virus-S Hemagglutinin Esterase (Non-Binding)

The coding sequence of MHV-S Hemagglutinin Esterase inactive (S45A) (1218bp) fused to human IgG-Fc followed by a 6xHis tag in the mammalian expression vector pcDNA3.1(-) (Invitrogen) was kindly provided by Colin Parrish (MHV-S-HE_Fc_His6) and described elsewhere. The Hisx6 and associated stop codon was deleted by Q5 site directed mutagenesis (New England Biolabs). Two complimentary oligonucleotides corresponding to the Avi-tag (Avidity) sequence GGTCTGAACGACATCTTCGAGGCTCAGAAAATCGAATGGCACGAA incorporating Hindlll overlapping ends were synthesized (Integrated DNA Technologies) and annealed at 100°C for 5 mins followed by slow cooling. The annealed oligos were cloned into the Hindlll cut site of MHV-S-HE_Fc to produce MHV-S_HE_Fc_Avitag. Non-binding mutant (F212A) was generated from template MHV-S_HE_Fc_Avitag using the Q5 Site-Directed Mutagenesis Kit (New England BioLabs)

#### SGRP-7 – Bovine Coronavirus-Hemagglutinin Esterase (Inactive); Bovine Corona virus-Hemagglutinin Esterase (non-binding)

The BCoV-Mebus HE ORF was amplified by PCR in junction overlap with a PCR amplification of the pFastBac1 clone of PToV-P4 He-Fc/Avi (see SGRP-9), but minus PToV HE sequence. Gibson assembly allowed for in-frame addition of BCoV-Mebus HE into the construct in direct replacement of PToV-P4 HE. This strategy maintained all vector, signal, and Fc-fusion/Avi architecture between constructs. Non-binding mutant F211A was generated by Q5 site-directed mutagenesis (New England Biolabs)

#### SGRP-8 – Clostridium tetani Heavy Chain/B Subunit; Clostridium tetani Heavy Chain/B Subunit (non-binding) mutant R1226L/W1289A; Clostridium tetani Heavy Chain/B Subunit (non-binding) mutant W1289A

The optimal receptor-binding domain (OBD) (100, 100) of Tetanus Toxin (Heavy Chain/B Subunit) was synthesized by GENEWIZ (incorporating flanking 5’ Hindlll and 3’ Nco1 restriction sites) and codon optimized for expression in E.Coli (a.a 1105>1315). pGEX-3X containing an irrelevant gene (IG) was engineered to contain a SNAPf_Hisx6 tag in the EcoR1 restriction site for use as an intermediate template (pGEX-3X_IG_SNAPf_Hisx6) and described elsewhere. Briefly, a Hisx6 sequence tag was inserted into the BamH1 and Xho1 sites of MCS2 of pSNAPf (New England Biolabs) to create pSNAPf_Hisx6. SNAPf_6xHis was amplified from this construct by PCR using VENT Polymerase (NEB), forward primer 5’-AAAA<u>GAATTC</u>AACCATGGACAAAGACTGCGAAA-3’ and reverse primer 5’-TTTT<u>GAATTC</u>ATTAGTGATGGTGATGGTGATGGGA-3’ (EcoR1 site are underlined). PCR products were digested with EcoR1 and sub-cloned into the EcoR1 site of the pGEX-3X_IG). pGEX-3X_IG_SNAPf_Hisx6 was then cut with Hindlll and Nco1 to directly replace IG with Hindlll and Nco1 cut B Subunit (OBD). Non-binding double mutant R1226L/W1289A and non-binding single mutant W1289A were generated by Q5 site-directed mutagenesis (New England Biolabs). Constructs were sequence verified.

#### SGRP-9 - Porcine Torovirus Hemagglutinin Esterase (inactive);Porcine Torovirus Hemagglutinin Esterase (non-binding)

The coding sequence of PToV Hemagglutinin Esterase inactive (S46A) and PToV Hemagglutinin Esterase non-binding (F271A) (1176bp) fused to human IgG-Fc followed by a 6xHis tag in the Insect expression vector pFastBac1 (ThermoFisher) was kindly provided by Colin Parrish (PToV-HE_S46A-Fc_Hisx6/pFastBac1; PToV-HE_F271A-Fc_Hisx6/pFastBac1) and described elsewhere. Both PToV-HE_S46A-Fc_Hisx6 and PToV-HE_F271A-Fc_Hisx6 were sub-cloned into pAC6 (Avidity) in-frame with the 3’Avitag (Avidity) sequence GGTCTGAACGACATCTTCGAGGCTCAGAAAATCGAATGGCACGAA. The resulting Avi-tagged constructs were amplified by PCR using pfu polymerase and the primers 5’ ATATA<u>GCGGCCGC</u>TTTCTCGAGGCATAAGGAACACAC; Not1 site underlined and 3’ CGCGC<u>GGTACC</u>TTATTCGTGCCATTCGATTTTCTGAG; Kpn1 site underlined. PCR products were digested with Not1 and Kpn1 and sub-cloned into the corresponding sites of PToV-HE_S46A-Fc_Hisx6/pFastBac1 and PToV-HE_F271A-Fc_Hisx6/pFastBac1, which were cut with Not1 and Kpn1 thus removing the original untagged sequences. The final constructs were sequence verified.

#### Y.Pestis SubB

The full-length coding region of YPSubB (420 bp) incorporating a 3’ Hisx6 tag in the bacterial expression vector pBAD18 was kindly provided by James Paton and is described elsewhere.

#### GspBBR; GSpBBR (non-binding); UB10712BR; UB10712BR (non-binding)

pGEX3X-GspBBRΔcnaA (containing just the Siglec and Unique domains of GspBBR) and pGEX3x-10712BR were constructed previously (2016 Glycobiology). The SNAPf_Hisx6 coding sequence was inserted in-frame into the EcoRI site at the 3’end of the BR sequences as described for SGRP-3 above. The non-binding mutant BR-SNAPf_Hisx6 sequences were synthesized (LifeTechnologies) and then used to replace the BR segment (HindIII-NcoI) of pGEX3X-GspBBRΔcnaA- SNAPf_Hisx6.

#### E. coli SubB; E. coli SubB (Jennings mutant); E.coli SubB (non-binding)

The coding region of E. coli SubB (423 bp) in pET-23 (+) bacterial expression vector was provided by Michael Jennings. Deletion of S106 and T107 was performed from the wt template above using the Q5 Site-Directed Mutagenesis Kit (New England BioLabs) and sequence verified. Mutant S106;T107 was cloned into pET28b which had been modified to contain a 3’ SNAPf_Hisx6 tag. This was subsequently engineered to contain an S12A mutation (non-binding) using Q5 mutagenesis.

#### CD22-Fc_ACP

A CD22 cDNA fragment encoding the first two Ig-like domains fused to an EK-hIgG-Fc fragment was amplified by PCR and cloned into the mammalian expression vector pACP-tag(m)-2 (New England Biolabs).

### Expression and purification of proteins in bacterial systems

#### Overexpression and purification of YenB and its nonbinding mutant

A single colony of BL21(DE3) competent cells (NEB, C2527H), transformed with YenB (or its mutant YenB S31A/Y100F) was inoculated in 25 ml of LB media in presence of 100 μg/ml ampicillin (Millipore, 171254-25GM) and grown overnight at 37 °C with mild shaking of 225 rpm. 20 ml of this primary culture was added to 1 liter of LB media supplemented with 100 μg/ml ampicillin and grown at 37 °C until optical density at 600nm reached to 0.2–0.3. To induced protein overexpression, arabinose (Sigma, A3256-25G) was added to the culture to make final concentrations 0.2% v/v, and the culture was incubated at 25 °C for 5–6 hours with mild shaking of 225 rpm. Bacteria was pelleted at 5000 g, 20 min, 4 °C and suspended in resuspension buffer (50 mM Tris-HCl, pH 8, 300 mM NaCl, 2 mg/ml lysozyme (Sigma, L6876), 1 mM EDTA, 10% glycerol in presence of DNAse (Sigma, D4263-5VL) and Protease inhibitors (Millipore, 5393134) at 4 g/ml. Pellet was resuspended, sonicated (30secs-ON/OFF, 10 cycles) and centrifuged (10000 g, 4 °C, 30 min) to exclude insoluble fractions. The soluble fraction was resuspended with preequilibrated Ni-NTA resin (Qiagen, 1018600) and incubated for 1 hr at 4 °C with end-to-end rotation. This slurry was passed through purification column (Biorad, PolyPrep 731-1550) and washed with 5 ml of 50 mM Tris-HCl, 300 mM NaCl, pH 8, followed by 2 washes with 5 ml of 50 mM Tris-HCl, 300 mM NaCl, pH 8, 30 mM Imidazole (Sigma, I2399-100G). The protein was eluted in 7 aliquots of 700 μl each with elution buffer (50 mM Tris-HCl, 100 mM NaCl, 300 mM Imidazole supplemented with protease inhibitors) and analyzed by SDS-PAGE. Aliquots with optimal protein concentrations were collected and transferred to PBS (Millipore, Amicon ultra centrifuge filter), supplemented with protease inhibitors, quantified (BCA, Pierce, Thermo Scientific 23225) and saved at −80°C until further steps.

#### Overexpression and purification of PltB and its nonbinding mutant

A single colony of BL21DE competent cells (NEB, C2527H), transformed with PltB (or its mutant-PltB S35A) was inoculated in 25 ml of LB media in presence of 50 μg/ml kanamycin (Sigma, K1377-5G) and grown overnight at 37 °C with mild shaking of 225 rpm. 10 ml of this primary culture was added to 0.5lit of LB media (Millipire, 1.10285.0500) supplemented with 50 μg/ml kanamycin and grown at 37 °C until optical density at 600 nm reached to 0.6–0.7. PltB overexpression was induced by 0.5 mM IPTG (Apex, 20-109) at 29 °C for 12–14 hours with shaking of 250 rpm. Bacteria was pelleted at 5000 g, 20 min, 4 °C and suspended in resuspension buffer (50 mM Tris-HCl, pH 8, 150 mM NaCl, 2 mg/ml lysozyme, 1 mM EDTA, DNAse and protease inhibitors) at 4 g/ml. Pellet was resuspended, sonicated (30 secs ON/OFF, 10 cycles, Fischer Scientific, 550 Sonic Dismembrator) and centrifuged (10000 g, 4 °C, 30 min) to exclude insoluble fractions (Sorvall, RC6 plus, Thermo). The soluble fraction was resuspended with preequilibrated Ni-NTA resin (Qiagen, 1018600) and incubated for 1 hr at 4 °C with end to end rotation. This slurry was passed through purification column (Biorad, PolyPrep 731-1550) and washed with 5 ml of 50 mM Tris-HCl, 150 mM NaCl, pH 8, followed by 2 washes with 5 ml of 50 mM Tris-HCl, 300 mM NaCl, pH 8, 50 mM Imidazole (Sigma, I2399-100G). The protein was eluted in 7 aliquots of 700 μl each with elution buffer (50 mM Tris-HCl, 100 mM NaCl, 300 mM Imidazole supplemented with protease inhibitors) and analyzed by SDS-PAGE. Aliquots with optimal protein concentrations were collected and transferred to PBS, supplemented with protease inhibitors, quantified (BCA, Pierce, Thermo Scientific 23225) and saved at −80 °C until further steps.

### Overexpression and purification of HsaBR and its nonbinding mutant

A single colony of BL21(DE3) competent cells, transformed with HsaBR (or its mutant HsaBR R340E) was inoculated in 25 ml of LB media in presence of 100 μg/ml ampicillin (Millipore, 171254-25GM) and grown overnight at 37^°^C with mild shaking of 225 rpm. 10 ml of this primary culture was added to 0.5lit of LB media supplemented with 100μg/ml ampicillin and grown at 37 °C until optical density at 600 nm reached to 0.7–0.8. Protein overexpression was induced by 0.5 mM IPTG (Apex, 20-109) at 24 °C for 12–14 hours (or 37 °C, 3 hrs) with shaking of 250 rpm. Bacteria was pelleted at 5000 g, 20 min, 4 °C and suspended in resuspension buffer (50 mM Tris-HCl, pH 7.5, 150 mM NaCl, 2 mg/ml lysozyme, 1% Triton X-100, 1 mM EDTA, DNAse and protease inhibitors) at 4 g/ml. Pellet was resuspended, sonicated (30 secs ON/OFF, 8 cycles) and centrifuged (10000 g, 4 °C, 30 min) to exclude insoluble fractions. The soluble fraction was resuspended with preequilibrated GST Sepharose resin (Thermo, 17-0756-01) and incubated for 1 hr at 4 °C with end to end rotation. This slurry was passed through purification column (Biorad, PolyPrep 731-1550) and washed with 10 bed volumes of resin by 50 mM Tris-HCl, pH 7.5, 150 mM NaCl, 1 mM PMSF, 0.5% Triton X-100 twice, followed by 2 washes with 10-bed-volume with 50 mM Tris-HCl, pH 7.5, 150 mM NaCl, 1 mM PMSF. The protein was eluted in 7 aliquots of 700 μl each with elution buffer (10 mM reduced glutathione (Sigma 4251-5G in 50 mM Tris-HCl, pH 8, 150 mM NaCl, supplemented with protease inhibitors) and analyzed by SDS-PAGE. Aliquots with optimal protein concentrations were collected and transferred to PBS, supplemented with protease inhibitors, quantified by BCA and saved at −80 °C until further steps. Similar procedures were followed for expression and purification of GspBBR, UB10712BR and their nonbinding mutants.

#### Overexpression and purification of TeT-HCR, TeT-HCR 1226L, TeT-HCR W1289A and its nonbinding mutant

The procedure to purify TeT-HCR and its variant proteins remained comparable to protocol adopted for HsaBR, except for the IPTG induction conditions. For TeT-HCR, proteins were overexpressed at 26 °C for 4 hr with shaking at 250rpm.

### Expressions and purification of proteins in mammalian expression system

#### Overexpression and purification of MHV-HE-Fc and hCD22_Fc

HEK293 cells were transiently transfected by endotoxin free construct of MHV-Fc virolectin or its mutant in presence of polyethyleneimine (PEI, Polyscience Inc, 23966). Precisely, 18 μg DNA was vortexed with 54 μl of 1 mg/ml PEI in 2.4 ml of advanced DMEM (no serum, Gibco-12491-015) and added to each of 15cm dish of 80–90% confluent cell in 20 ml (advanced DMEM, 2%FCS, 1X P/S, 1X glutamine) and incubated 24 hr at 37 °C in incubator. Post incubation, the media was replaced by 20 ml basal media (96 ml RPMI 1640 (Invitrogen 11875-135), 96 ml DMEM (Invitrogen 12430-104), 2 ml of antibiotics 100X, 2 ml 200 mM L-glutamine, 2 ml 100 × Nutridoma (Nalgene, Nutridoma-SP, 11011375001), 2 ml of 100 mM sodium pyruvate) per plate and incubated for 4 days. The media was gently collected and centrifuged at 1000 g, 15 min, 4 °C to remove cell debris. The supernatant was adjusted to 10 mM Tris-HCl, pH 8, filtered with a 0.2 μM filter unit (Corning, 430320) and incubated with 0.5 ml protein-A-sepharose (nProtein A Sepharose 4 Fast flow, GE healthcare #1705280) at 4 °C for 24 hr. Resin was collected in purification column, washed with 10 ml of 10 mM Tris-HCl, pH 8, 150 mM NaCl twice and then resin was resuspended in 2 ml of 20 mM HEPES pH 7. Resin was treated with 25mU of Arthrobacter ureafaciens sialidase (EY laboratory, EC-32118-5) at room temperature for 1 hr and then washed with 10 ml of 10 mM Tris-HCl, pH 8, 150 mM NaCl to remove sialidase. MHV-Fc protein was eluted with 2 ml of 0.1 M glycine pH 3.0 in a tube, already containing 0.3 ml of cold 1 M Tris-HCl, pH 8, to neutralize pH. The elutant was buffer exchanged to PBS, supplemented with protease inhibitors, quantified by BCA assay and saved at −80 °C until further steps. Similar procedure was followed for expression and purification of hCD22-Fc.

### Expressions and purification of proteins in insect expression system

Clones of pFastBac1 constructs were used to generate recombinant bacmids in DH10Bac following the manufacturers protocol (ThermoFisher). Recombinant baculoviruses were recovered by transfection of the bacmids into Sf9 insect cells using TransIt-Insect reagent (Mirus). Viruses were then used to infect suspension High Five cells grown in Express5 media and supernatants harvested ∼60–72 hours post-infection. Proteins were purified from the infected cell culture supernatants by binding to a HiTrap ProteinG HP column (GE Healthcare Life Sciences, Piscataway, NJ) and eluted with 0.1 M citrate, pH 3.0 (pH neutralization to 7.8 with 1 M Tris-HCl, pH 9.0) using ÄKTA FPLC system (GE Healthcare Life Sciences). The HE-Fc containing fractions were dialyzed in PBS and concentrated using 30kD Amicon Ultra-15 filters (EMD Millipore). Purified proteins were observed by SDS gel migration as well as anti-human IgG Fc probing by Western. Protein concentrations were determined from A280 readings and MicroBCA Assay. Proteins were primarily aliquoted and stored at −80 °C.

#### Biotinylation of Proteins

Biotinylation of SNAP tagged proteins and Avi-tagged proteins were performed as suggested by manufactures (NEB, Avidity). Direct biotinylation of proteins for example-YenB, PltB and Neu5Gc-IgY was achieved by NHS mediated biotin tagging at 20:1 molar ratio as per manufacturer’s protocol (Thermo scientific, EZ-Link Sulfo-NHS-LC Biotin, Cat#21326). Biotinylated proteins were concentrated, and buffer exchanged to PBS with microcon centrifugation columns according to the molecular weights of proteins. Finally, elutions were estimated for protein concentration and stored in −80 °C freezer. In addition to the functional assays on sialoglycan microarray and sera ELISA experiments, high efficiency of biotinylation was also confirmed by a pull-down assay using streptavidin beads (Thermo, 20347). Briefly, proteins biotinylated and mock biotinylated were tested for their affinity and subsequently elution from streptavidin beads. Pulled down biotinylated proteins were also compared parallelly to the initial concentration of protein to observe the approximate fractions of proteins acquiring biotin in biotinylation tagging procedures.

#### Sialoglycan microarray with SGRPs

As described by accompanying article by *Sasmal et al*.

#### Sialic acid release, DMB derivatization, and HPLC analysis

Bound Sias from serum samples were released using 2 M acetic acid hydrolysis, 80 °C for 3 hour. The hydrolyzed samples were passed through a Microcon-10 filter (Sigma-Millipore); the released Sias were recovered in the filtrate. The Sias were derivatized with DMB (112). Briefly, 7 mM DMB, 1.4 M acetic acid, 0.75 M β-mercaptoethanol, 18 mM sodium hydrosulfite, at 50 °C for 2.5 hour, in the dark. The DMB-Sias were separated on a Gemini C18 column, 4.6 × 250 mm, 5 µ (Phenomenex; Torrance, CA). Isocratic elution was employed using 7% methanol, 8% acetonitrile, 85% water for 50 min at 0.9 ml/min flow. The eluant was monitored by fluorescence with Ex373 and Em448. Sialic acid quantitation was done by derivatizing commercial Neu5Ac (Nacalai, USA; San Diego, CA) and comparing the area under the peak of known amounts.

#### Serum Binding assay with SGRPs

A saturable amount of sera (equivalent to 50μg proteins), reconstituted in PBST (0.1% Tween 20) were added to individual wells in ELISA plates (Corning, 9018) and kept overnight at 4°C for adherence. Unattached sera were removed, and wells were briefly rinsed with PBST before addition of 200 μl, 0.5% cold fish gelatin (Sigma-G7765-1L, diluted in PBST) for blocking. Lectins, SGRPs or other probes were reconstituted in diluting buffer (PBST with 0.05% cold fish gelatin) at desired concentrations and added to serum coated wells for 1hr (or otherwise specified) at room temperature. Specifically, for MAL-I (VECTOR LAB, B1315) and MAL-II (B1265), 10 mM CaCl2 and 10 mM MnCl2 were added to diluting buffer to maximize lectin binding. Post incubation, wells were vigorously washed with 250μl PBST, 5 times and then incubated with Avidin-HRP (Biolegend, 405103) for biotinylated probes at 1:2000 dilution for 45 minutes at room temperature. Wells were again washed vigorously washed with 250 μl PBST for 6–7 times. 100 μl of TMB (BD OptEIA-55214) was added to each well as substrate for HRP’s enzymatic activities and incubated for color development. The reactions were stopped by adding 40 μl of 2 N H2SO4 after 30 minutes, or if the wells specified as negative controls begin to develop color. Plates were read at 450 nm and readings were processed to observe binding patterns of tested molecules (Perkin Elmer, Enspire Alpha). For every experiment, wells treated with avidin-HRP only served as negative control of binding, along with non-binding mutants of probes, whenever applicable.

For serum binding experiment with nonbiotinylated proteins, binding of Fc-tagged proteins was probed by whereas binding of His6-conjugated proteins was probed using anti-His antibody (mouse, Gene Script, #A00186, 1:4000) and subsequently with anti-mouse-HRP (CTS, #7076S, 1:1500). In certain probes discussed in article (for example, LFA (EY Lab, BA1501-1, CHE-FcD etc.), the pre-conjugated combination of probe and secondary antibody was adopted for stronger signals, as mentioned in corresponding figure legends.

#### Mild Periodate and Base treatment of sera

To remove *O*-acetyl esters and render all Sias sensitive to mild periodate treatment, serum-coated ELISA plates were expose to strong NH4OH vapors for 24 hrs at room temperature. In parallel, another plate coated with sera were incubated in humidified chamber filled with triple distilled water at room temperature as control set of base treated sera. Wells were washed 3 times with PBS briefly to remove any residual NH4Cl. Subsequently, plates were washed three times with PBS pH 6.5 to pre-equilibrate for mild periodate treatment, if required. To ascribe Sia specificity, serum-coated ELISA plates were treated with mild periodate to selectively cleave the Sia side-chain. For sodium periodate treatment in base treated/untreated plate, wells were incubated with 200μl of freshly prepared cold 2 mM sodium metaperiodate (Fischer, S398-100) in PBS, pH 6.5 in dark, 4°C, precisely for 20min. The reaction was stopped by the addition of 50 µl of 100 mM sodium borohydride (Fischer, S678-25) in PBS, pH 6.5 (final concentration of 20 mM), followed by a 10-min incubation at room temperature with gentle shaking (the borohydride inactivates the periodate). Concurrently, as a mock control, periodate and borohydride solutions were premixed (4:1), and wells were incubated with 250 µl/well side by side with the periodate-treated wells. To remove resulting borates, wells were then washed three times (10 min each wash) with 50 mM sodium acetate, pH 5.5, containing 100 mM NaCl, followed by three washes with PBS, pH 7.4. Base and/or mild periodate treated sera were used to determine Sia specific serum binding, or OAc ester specific Sia binding in SGRPs or lectins. (113)

#### Western Blotting

Commercial animal sera (rat, equine, goat, guinea pig, rabbit, sheep; Sigma, St.Louis, MO), and wt mouse and human serum (Valley Biomedical, Winchester, VA) and in house *Cmah*^−/−^ mouse serum were chromatographed on 12% CosmoPAGE Bis-Tris precast gels (Nacalai, USA, NU00212). The gels were run in CosmoPAGE MOPS running buffer pH 7 (Nacalai, USA, NU01004), with ice packs to keep running buffer temperature below 23 °C. The proteins were transferred to Immobilon FL (EMD Millipore Corp IPFL00010) membrane in transfer buffer containing 10% methanol and 0.2% SDS. Membranes were blocked overnight at 4 °C with 0.5% Cold water fish skin gelatin in PBST (PBS pH 7.4 with 0.1% Tween 20), with gentle agitation. The membranes were incubated at room temperature for 2 hrs with 1 µg/ml with all but SGRP5*^IgY^*, which was used at 0.33 µg/ml, diluted with blocking buffer without Tween. Also, SGRP4*^MHV^* and SGRP9*^PToV^* were used at 2 µg/ml. Amounts of serum proteins immunoblotted for each SGRP were adjusted to highlight their ligand preferences. Specifically, 25 µg protein per well was loaded for SGRP1*^YenB^*, SGRP2*^PltB^* and their nonbinding variants, 50µg protein per well was loaded for SGRP3*^Hsa^*, SGRP4*^MHV^*, SGRP7*^BCoV^*, SGRP9*^PToV^* and their nonbinding mutants, while 12.5 µg protein per well was loaded for SGRP5*^IgY^*, SGRP5*^IgY^*NB and SGRP6*^SNA^* (Vector lab, B1305). Secondary antibody incubation was with Streptavidin IRDye 680 (Licor, 925-68079) at 1:10,000 dilution in PBST at room temperature. Signal was read with Odyssey infrared imager (Licor Biosciences, Lincoln, NE). Interrogating SGRPs and corresponding non-binding SGRPs were used at the same concentration and all images were obtained at the same time using the same imager settings.

#### Histology Procedure

Frozen sections of various organs from the wild type and *Cmah*^−/−^ mouse were air dried for 30 minutes, rehydrated in Tris-Buffered Saline pH 7.5 and overlaid with 0.5% Fish gelatin, which was then tipped off for blocking the endogenous biotin suing the Avidin Biotin blocking kit from Vector labs (SP-2001). Sections were then fixed for 30 minutes with 10% Neutral buffered formalin (VWR-89370-094), washed and overlaid with either the binding or non-binding probes, each used 5 micrograms/milliliter. Separate slides (Globe, 1358W) were then overlaid with each of the Biotinylated probes and incubated in a humid chamber at room temperature for 30 minutes, followed by washing in TBS and then overlaid with Cy3 Streptavidin (Red, Jackson Laboratory, 016-160-084) for 30 minutes, washed and then nuclei were counterstained with Hoechst (Blue, Molecular Probes, H3570) for two minutes, before washing and cover-slipping using an aqueous mounting medium (Vector lab H5501). Digital images were captured using the Keyence microscope, and organized using Adobe Photoshop.

#### Flow Cytometry

SGRPs and probes were probed at 5 μg/ml (or otherwise, mentioned in figure legends) for their RBC binding properties. For this experiment, blood from several animal species was resuspended in PBS (4 °C) to make 2% volume/volume dilution. 200 μl of this blood suspension was supplemented with 200 μl of 10 μg/ml probe dilution in PBS (4 °C) in a tube to make final blood suspension and probe dilutions at 1% and 5 μg/ml (unless mentioned otherwise). 400μl of this blood suspension was incubated ice for 1hr with intermittent gentle mixing to avoid settlement of RBCs. After incubation, 1 ml ice cold PBS was added to each tube and centrifuged at 1000 RPM, 3 minutes at 4 °C. The washing was repeated 2 more times, and then pellet was gently broken into 300 μl of 1 μg/ml dilution of Streptavidin Cy3 (Jackson 016-170-084) and incubated for 45minutes in ice. For autofluorescence control of RBCs, blood suspension was never treated either by probe or fluorescence antibody, while for fluorescent secondary antibody control, a mock treated blood was incubated with equivalent streptavidin-Cy3 dilution in similar conditions. After incubations, all tubes were added with 1 ml ice cold PBS and washed 3 times with 1 ml PBS. Final pellet was gently broken in 500 μl of PBS and fluorescence was analyzed in FL2 channel in FACS Caliber flow cytometer. The settings for acquisition of data and analysis remained same for all probes and their nonbinding mutants in an experiment.

## Acknowledgments

This work was supported by NIH Grants U01CA199792 and R01GM32373 (to A.V.), and additional generous financial support from BioLegend, San Diego.

## Abbreviations

SGRP: Sialoglycan Recognition Probe
Sia: Sialic Acids
Siglec: Sialic acid-binding immunoglobulin-type lectins
YenB: Yersinia enterocolitica, toxin subunit B
HE: Hemagglutinin Esterase
HA: Hemagglutinins Adhesins
Neu5Ac: N-Acetylneuraminic acid
Neu5Gc: N-Glycolylneuraminic acid

